# Universal approach to wave-optical calculations of point spread functions in microscopy (and beyond)

**DOI:** 10.64898/2026.04.28.721333

**Authors:** Ivan Gligonov, Lars Loetgering, Francisco Tenopala-Carmona, Chia-Lung Hsieh, Ingo Gregor, Jörg Enderlein

**Affiliations:** Third Institute of Physics (Biophysics), Georg August University, Friedrich-Hund-Platz 1, 37077 Göttingen, Germany; Carl Zeiss Microscopy GmbH, Carl Zeiss Promenade 10, D-07745 Jena, Germany; Humboldt Centre for Nano- and Biophotonics, Institute of Light and Mater, Department of Chemistry, University of Cologne, Greinstr. 4-6, D-50939 Cologne, Germany; Institute of Atomic and Molecular Sciences, Academia Sinica, Taipei, Taiwan; Cluster of Excellence "Multiscale Bioimaging: from Molecular Machines to Networks of Excitable Cells” (MBExC), Universitätsmedizin Göttingen, Robert-Koch-Str. 40, Göttingen, 37075, Germany

## Abstract

Optical microscopy is fundamental to modern life-science research, yet interpreting its results requires precise modelling of point spread functions (PSFs) within complex environments. This manuscript introduces a versatile and efficient approach to wave-optical PSF calculations that extends existing frameworks by incorporating detection PSF modelling through the principle of reciprocity. Accompanying this work is a free MATLAB software package centred on a single, minimalistic core function, **PlaneWaveExc.m**, which utilizes a plane-wave superposition based on the Richards-Wolf model. Despite its simplicity, the framework accounts for „real-life” complexities such as systemic aberrations, arbitrary amplitude and phase modulations, and stratified media with complex-valued refractive indices. We demonstrate the software’s broad applicability through diverse case studies, including single-molecule imaging, STED microscopy, the segmented aperture of the James Webb Space Telescope, and coherent wide-field iSCAT microscopy. Each example is supported by dedicated scripts to facilitate adaptation for specific research needs.

## 1. Introduction

Optical microscopy remains a cornerstone of life-science research. Recent years have seen a transformative surge in methods aimed at enhancing resolution, contrast, and information density. To accurately interpret these images and model experimental outcomes, precise calculations of the PSF—the fundamental response of an imaging system to a point source—are indispensable. In particular for (3D) image deconvolution, an accurate knowledge of the PSF is of paramount importance, see e.g. refs. [1,2]. While several software packages currently exist to address this need, they must balance mathematical rigor with speed, user accessibility, and the versatility required to model complex optical environments.

A substantial body of literature and various software tools have focused on the acceleration of Point Spread Function (PSF) calculations, most notably in references [3–7]. Especially the work by Leutenegger et al. [3] is widely considered the gold standard for calculating focus excitation fields. However, calculating the *detection PSF*—defined as the efficiency with which an emitter’s signal is captured as a function of its spatial position—is typically far more complex.

Conventionally this process begins by expanding the electric field of an emitter (typically modelled as an oscillating electric dipole) into a superposition of plane waves using Weyl or angular spectrum representations [8,9]. These waves must then be individually traced through the entire optical system to derive the final Poynting energy flux at the detector plane. While this approach is straightforward in a homogeneous medium, the complexity scales significantly when the emitter is embedded within a layered structure. In such environments, one must account for field modifications—including reflections, transmissions, and interference—within the layers before the field can even be propagated through the imaging optics. This computational burden is further compounded when dealing with birefringent materials, which introduce highly non-trivial anisotropic field distributions, or when modelling higher-order sources such as electric quadrupoles or magnetic dipoles. In these scenarios, the forward calculation of the detection PSF is substantially more involved than that of the excitation PSF.

In this manuscript, we extend the existing framework by calculating the detection PSF through the *principle of reciprocity* [10–14]. This principle asserts that the physics of an emitter producing a photon that is subsequently captured by a point detector is reciprocal to a scenario where the detector ‘emits’ a photon that is absorbed by the emitter. By employing this symmetry, we can utilize the exact same computational routine used for focus excitation fields to determine the detection efficiency of an emitter. Beyond being conceptually more elegant than full emission-field modelling, our approach offers a major practical advantage: it requires only a single, unified computational framework to derive both the excitation and detection PSFs, regardless of the complexity of the sample environment.

Accompanying this paper is a free MATLAB (The MathWorks, Inc.) software package centred on a single, minimalistic core function. Despite its simplicity, this function is capable of modelling a diverse array of imaging modalities. We have placed special emphasis on „real-life” complexity, including: (i) Modelling realistic optical imperfections; (ii) Handling both amplitude and phase modulations in excitation and detection; and (iii) Support for stratified media with arbitrary, complex-valued refractive indices.

In the following chapter, we provide a brief overview of the physical principles underlying our PSF calculations and detail the architecture of the core MATLAB function. The third chapter provides several concrete applications of the software. To assist the reader in adapting the code for specific research requirements, each example is accompanied by a dedicated MATLAB script that reproduces the figures presented in this work.

The case studies include:

1. Stratified Samples: Calculating focal distributions in multi-layered media.
2. Wide-field Microscopy: Incorporating the effects of optical aberrations.
3. Confocal Detection: Analysing the impact of pinhole misalignment.
4. Single-Molecule Imaging: Modelling defocused imaging of single emitters.
5. Scanning Microscopy: Imaging single molecules with complex excitation polarization.
6. STED Microscopy: Modelling Stimulated Emission Depletion imaging.
7. The James Webb Space Telescope: A case study for non-trivial apertures.
8. Phase Modulation: Demonstrating the Double-Helix PSF.
9. Nonlinear Imaging: Modelling Super-resolution Optical Fluctuation Imaging (SOFI).
10. Coherent Imaging: Modelling wide-field Interferometric Scattering (iSCAT) microscopy.

## 2. Theoretical background

### 2.1 Focusing of a plane wave through an objective

We start with focusing a linearly polarized plane wave through an objective as shown in Figure 1a. Following the classical works by Wolf [15] and Richards and Wolf [16], the electric field distribution in the focal region is given by the plane-wave superposition

**Figure 1.**
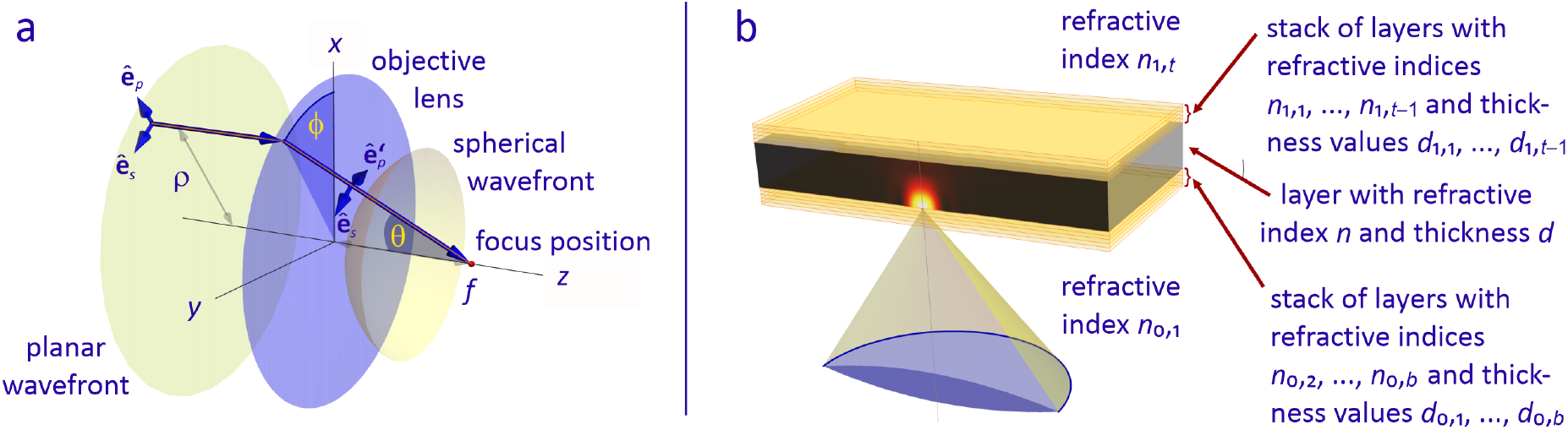
Panel a: Geometry of focusing a plane wave through an objective lens. Panel b: Geometry of a layered sample consisting of a bottom and top stack of planar layers with different refractive indices and thickness values enclosing the sample layer with refractive index *n* and thickness *d*.

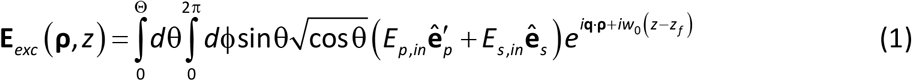

The integration variables are the polar angle θ and azimuthal angle ϕ as shown in Figure 1a. The lateral component **q** of the wave vector is given by

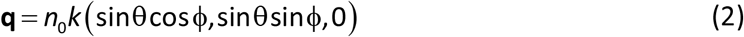

and the axial component *w*_0_ is given by

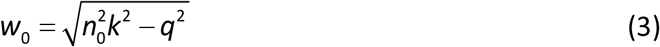

Here, *k* = 2π /λ is the length of the wave vector of a plane wave in vacuum, and *n*_0_ is the refractive index of the objective’s immersion medium. The unit vectors 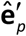 and **ê**_*s*_ point along the *p*and *s*-polarization of plane waves traveling along the direction (**q**,*w*_0_), and *z*_*f*_ is the focus position with respect to the coordinate origin at *z* = 0.

The *E*_*p,in*_ = cosϕ and *E*_*s,in*_ = sinϕ are the electric field amplitudes of the *p*- and *s*-wave components for focusing an incoming plane wave polarized along the *x*-axis (ϕ = 0) and having unity amplitude. The factor 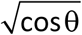 in the above expression ensures energy conservation upon focusing. The integration over θ extents to Θ, the maximum half angle of light collection of the objective. This angle is defined by the objective’s numerical aperture *NA* via *NA* = *n*_0_ sinΘ. The above equation holds true when focusing a plane wave into a homogeneous medium with a refractive index, *n*_0_, that matches the objective’s immersion medium. In many practical scenarios, however, the sample structure is significantly more complex, often consisting of multiple layers with varying refractive indices (as illustrated in Figure 1b). In such cases, the transmission and reflection of plane waves at these interfaces can substantially alter the amplitude and phase of the electric field at the focal plane. Consequently, the modified result is expresses as follows:

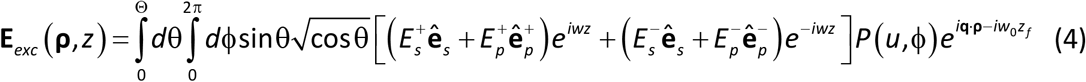

where the axial component *w* is given by

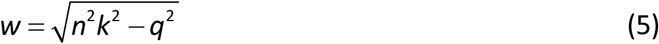

involving the refractive index *n* of the sample layer which can be significantly different from the *n*_0_ of the immersion medium. The superscripts + and – now refer to forward and backward propagating plane waves within the sample layer, generated by the manifold reflections from the stacks of layers below and above the sample layer. A description of how to find the corresponding electric field amplitudes 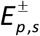 is detailed in the next section. The corresponding unit vectors 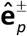 and **ê** in eq.(4) are defined by (see Figure 1b)

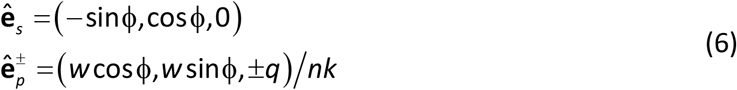

Eq. (4) also incorporates an additional pupil function *P* (*u*,ϕ) which accounts for arbitrary amplitude and phase modulations of the electric field incident onto the back focal plane (BFP). This function is defined in terms of the normalized pupil radius, *u* = sinθ sinΘ, and the azimuthal angle ϕ.

### 2.2 Propagation of plane waves through a stack of planar layers

It remains to determine the field amplitudes 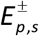 within the sample layer, accounting for the multiple transmissions and reflections of plane waves at the interfaces of a stratified stack with varying refractive indices. This will be done by using the method using the Transfer Matrix Method [17].

Consider the propagation of a plane wave with a lateral wave vector component **q** through a stack of *N* planar layers with refractive indices *n*_0_,*n*_1_,…,*n*_*N*+1_ and thickness values *d*_1_,*d*_2_,…,*d*_*N*_, where the refractive index *n*_0_ refers to the lower infinite half space and the refractive index *n*_*N*+1_ to the upper infinite half space. At each interface dividing layer *j* and layer *j* + 1, there are two plane waves with amplitudes 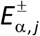 in layer *j* and two plane waves 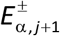 in layer *j* + 1, where the plus superscript refers to forward propagating waves with axial wave vector components 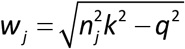, and the minus sign to a backward propagating waves with axial wave vector components - *w*_*j*_. The subscript α determines whether we consider a *p*-wave (α = *p*, polarization vector in plane of incidence) or an *s*-wave (α = *s*, polarization vector perpendicular to incidence plane). Taking into account the boundary condition that in-plane components of the electric and magnetic fields are continuous across interfaces, the electric field amplitudes across an interface are coupled by the equation

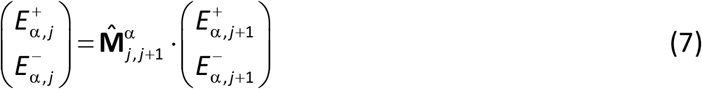

where the transfer matrix for *p*-waves is given by

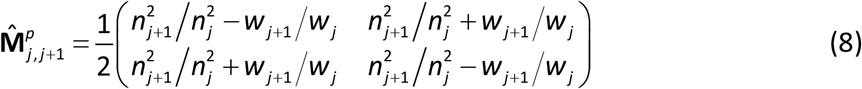

and for *s*‐waves by

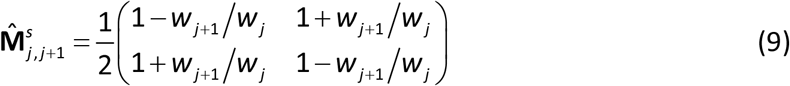

The propagation matrix within a layer, connecting the electric field amplitudes at interface (*j, j* + 1) with those at interface (*j* – 1, *j*) reads

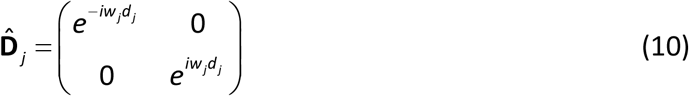

Consequently, for the whole stack we find

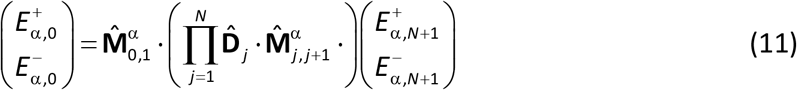

By setting 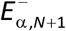 to 0 (no back propagating wave in the upper half space) and 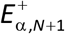 to 1, one can thus express 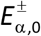 through 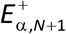. This allows for calculating the total reflection and transmission coefficients as

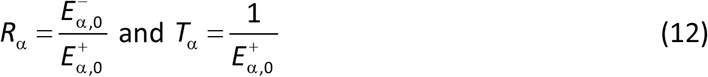

Consider a layer with refractive index *n* and thickness *d*, sandwiched between two stacks of layers as shown in Figure 1b. The compound transmission coefficient from bottom into the layer is denoted as *T*_*b*,α_. The compound reflection coefficients, measured from inside the layer towards the boundaries, are *R*_*b*,α_ for the bottom interface and *R*_*t*,α_ for the top interface. These coefficients are calculated according to the procedure described above.

For a plane wave incident from the bottom, the electric field amplitudes as used in eq. (4) are given by

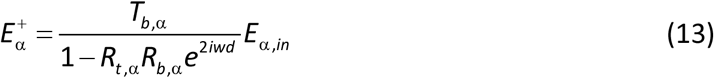

and

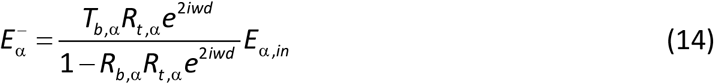

which accounts for the infinite number of multiple reflections between the top and bottom layer stacks.

### 2.3 The excitation PSF

Having calculated the electric field distribution **E**_*exc*_ (**ρ**, *z*) in the focal region, one can calculate the PSF of excitation (excitation PSF) for imaging a dipole emitter which has an absorption dipole amplitude vector **p**. At sufficiently low excitation intensities, the excitation rate of such a dipole emitter is proportional to the square of the scalar product of its absorption dipole moment **p** with the local electric field amplitude **E**_*exc*_ (**ρ**, *z*), so that the excitation PSF *U*_*exc*_ (**ρ**, *z*) is given by

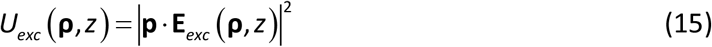

It should be noted that the excitation PSF is modified at high excitation intensities, which can pump molecules into intermediate photophysical states, such as the triplet state or photo-induced radical states. At very high intensities, the excited singlet state also becomes increasingly saturated, further altering the effective absorption rate. For a detailed discussion of such effect and their impact on the PSF, see e.g. ref. [18]. For the sake of simplicity, we do not consider these complex photophysical scenarios here, although our routines can be easily adapted to account for them.

In many cases, it can be assumed that imaging is performed with an isotropic distribution of emitter dipoles, in which case the PSF simplifies to

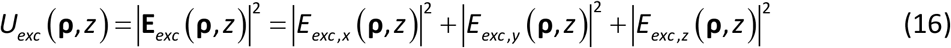

This assumption also holds for the common scenario where emitters rotate rapidly relative to the acquisition time, resulting in an isotropic time-averaged orientation.

### 2.4 The reciprocity principle and the detection PSF

We now turn to the detection PSF, which quantifies the efficiency with which a point source— typically a dipole emitter—is captured by a point detector at a specific coordinate in image space. The conventional „forward” approach to this problem begins with a plane-wave decomposition of the emitter’s electric field, such as the Weyl or angular spectrum representation [8,9]. These components are then traced through the optical system—a process mathematically analogous to the focusing model described in Section 2.1—to derive the final Poynting energy flux at the detector plane.

To circumvent these complexities, we present a significantly simpler approach that leverages our routines for modelling the excitation PSF. Our method is based on the principle of reciprocity [10–14]. Rather than calculating the probability of a photon emitted by the source reaching the detector, we analyse the inverse scenario: how effectively a photon „emitted” by a point detector would be absorbed by the emitter. By asserting that this absorption probability is equal to the detection probability, we bypass the need to compute unique field distributions for various emitter types or orientations.

Let us consider the electric field distribution in the sample space generated by a virtual point source—specifically, a detector pixel located on the optical axis in the image plane. We model this pixel as an ideal electric dipole oriented along the *x*-axis, which corresponds to *x*-polarized detection; the *y*-polarized case is subsequently obtained by a 90° rotation around the optical axis. This configuration is illustrated in Figure 2.

**Figure 2.**
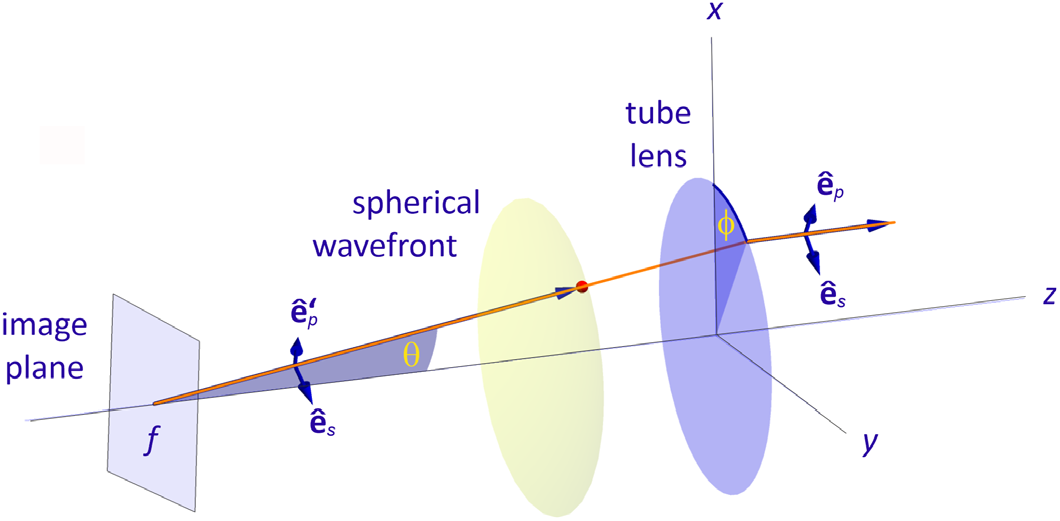
Illustration the path of a single light ray as it traverses the tube lens, which focuses the collected light onto the detector plane. To model this process using the principle of reciprocity, the calculation is inverted: we determine the electric field distribution generated by a virtual point dipole emitter located on the detector pixel for which the detection efficiency is being calculated.

To determine the electric field distribution in the BFP, we follow a procedure analogous to that in Section 2.1. The dipole moment vector is projected onto the unit vectors 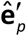 and **ê**_*s*_ (see Figure 2), yielding the corresponding electric field amplitudes in the BFP (tube lens plane in Figure 2). To satisfy energy conservation during the collimation by the tube lens, we include an apodization factor of 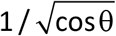 to account for energy conservation [15,16]. Assuming a dipole amplitude of unity, the field components are:

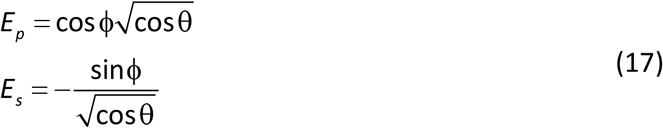

Consequently, the electric field amplitudes along the *x*- and *y*-directions in the BFP are given by

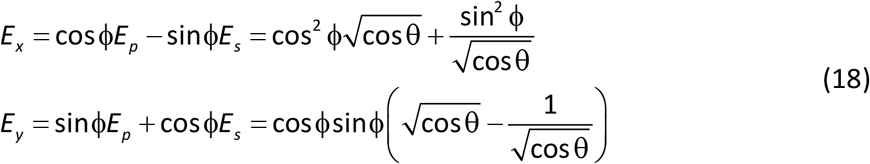

These equations demonstrate that the initially *x*-polarized electric field undergoes depolarization upon collimation by the tube lens. To estimate the magnitude of this effect, we analyse the maximum possible values of 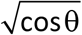 for an imaging system that uses an objective with numerical aperture *NA* and magnification *M*. According to Abbe’s sine law, we have sinΘ= *NA M*, where Θ is the maximum possible value of angle θ in image space. For a typical high-NA setup (e.g., a 1.2 NA water-immersion objective with magnification *M* = 60), the maximum angle in image space is θ_max_ = 0.0200 rad (1.15°). At such small angles, cosθ ≈ 1 to at least three decimal places, so that in eq. (18) one has *E*_*x*_ ≈ 1 and *E*_*y*_ ≈ 0. This indicates that the electric field in the BFP remains almost perfectly *x*-polarized, with negligible contributions along the *y*-direction. This approximation is even more robust when considering intensities, which scale with the absolute square of the amplitudes.

As a result, the electric field generated in the sample space by a detector pixel is, to a high degree of accuracy, identical to the field produced by the diffraction-limited focusing of a linearly polarized plane wave. Assuming the imaging system is *isoplanatic* (shift-invariant), we obtain the following identity for the detection PSF:

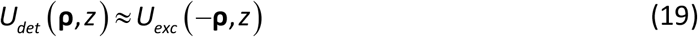

Note that the right-hand side must be evaluated at the emission wavelength rather than the excitation wavelength. While we utilize this approximation throughout this paper, we will later describe how to rigorously incorporate depolarization effects when discussing phase modulation of the back-focal wavefronts.

A critical detail involves the interpretation of the coordinates **ρ** and *z* in *U*_*det*_ (**ρ**, *z*). The lateral vector **ρ** represents a position on the image plane, *back-projected into sample space*. The axial coordinate *z* denotes the physical position of a point emitter along the optical axis that would produce the corresponding intensity distribution in the image plane.

Finally, for detection through a confocal aperture, the effective detection PSF *U*_*con*_ (**ρ**, *z*) is obtained by integrating *U*_*det*_ (**ρ**, *z*) over the aperture are *A*:

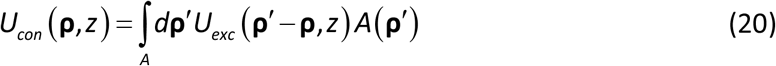

This calculation is most efficiently performed using the convolution theorem. Here, the aperture function *A*(**ρ**) represents the back-projection of the physical pinhole into the sample space—typically defined as unity within the clear aperture opening and zero elsewhere.

### 2.5 Core MATLAB function for PSF calculation

This manuscript is accompanied by a comprehensive, open-source MATLAB package for calculating and visualizing all PSFs described herein. The package is available on GitLab at https://gitlab.gwdg.de/ag_enderlein/universal_psf.

The numerical implementation of the focal field calculations is centred around a core MATLAB routine, **PlaneWaveExc**. This routine evaluates the double integral in eq.(4) in three steps. First, the algorithm computes the coefficients of the Fourier expansion of the pupil function with respect to the azimuthal angle ϕ. For each Fourier mode, the integration over ϕ is performed analytically. This step transforms the azimuthal dependence into a series of Bessel functions of the first kind, which are evaluated involving the polar angle θ. Finally, the remaining integral over the polar angle θ is computed numerically using Simpson’s rule.

**PlaneWaveExc** computes the complex electric field distribution in the focal region for an *x*-polarized plane wave focused by a high-NA objective. The routine is invoked as follows:

**exc = PlaneWaveExc(rhofield, zfield, NA, n0, n, n1, d0, d, d1, lambda, focpos, phirot, pupil, maxm, maxnum)**

It has the mandatory input variables **rhofield, zfield, NA, n0, n, n1, d0, d, d1, lambda, focpos**. The optional variables **phirot, pupil, maxm, maxnum** are discussed in detail below. The output **exc** is a MATLAB structure containing the calculated electric fields, the coordinates at which they were computed, and a record of all input variables.

The first two input variables determine the lateral and axial extent of the region of interest. The one-dimensional vector **zfield** defines the discrete axial positions (*z*_**k**_) where the electric fields are calculated. The variable **rhofield** can take two forms:

#### Radial vector

If **rhofield** is a one-dimensional vector, it defines the discrete radial positions (ρ_j_) from the optical axis. In this case, the calculated electric fields are then returned as Fourier components: **exc.fxc, exc.fxs, exc.fyc, exc.fys, exc.fzc** and **exc.fzs**. These represent the fields as a Fourier expansion in the angular variable ϕ (around the optical axis) as follows:

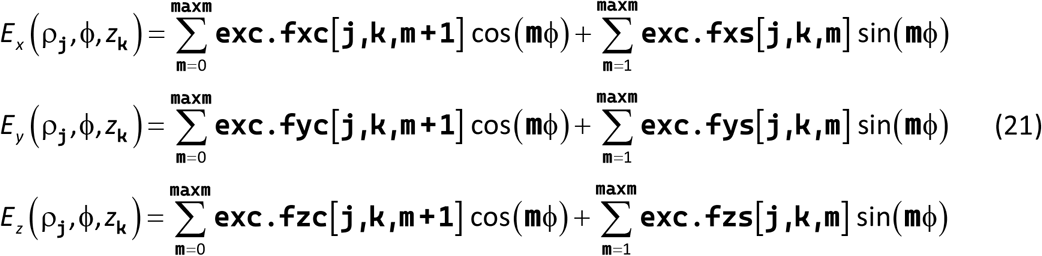

Here, ϕ is measured from the *x*-axis, and the ρ_**j**_ and *z*_**k**_ are the discrete positions defined in the vectors **rhofield** and **zfield**. The optional variable **maxm** defines the maximum number of Fourier components. For an ideal *x*-polarized plane wave, components exist only up to **maxm** = 2 (default value). For complex amplitude- or phase-modulated input fields, this number must be increased appropriately. This will be demonstrated in the next chapter, where we present modelling results for input fields with complex amplitude and phase modulations in the BFP.

#### Rectangular Grid

Alternatively, if **rhofield** is a structure with fields **rhofield.x** and **rhofield.y**, the routine computes the fields on a 3D rectangular grid generated by the MATLAB command **meshgrid(rhofield.x(:)’, rhofield.y(:), zfield)**. The structure **exc** will then contain the 3D matrices **exc.fieldx[j**,**k**,**l], exc.fieldy[j**,**k**,**l]**, and **exc.fieldz[j**,**k**,**l]**, corresponding to the grid positions *x*_**j**_, *y*_**k**_, and *z*_**l**_.

The input variable **NA** defines the numerical aperture of the objective used. The one-dimensional vectors **n0** and **n1** contain the refractive index values of all layers located below (**n0**) and above (**n1**) the sample layer, see Figure 1b. The vectors **d0** and **d1** represent the corresponding thickness values (in µm) for these layers. These vectors should have a length one less than **n0** and **n1**, respectively, to account for the semi-infinite outer layers. If **n0** or **n1** are single values, the space below or above the sample is assumed to be a homogeneous half-space with the specified refractive index. Consequently, if **n0** and/or **n1** are single numbers, then **d0** and/or **d1** must be set to an empty matrix []. The sample layer itself is characterized by refractive index **n** and thickness **d**, and is the region where the electric field calculations are performed. It should be noted that **n0** and **n1** can contain complex-valued numbers to account for layers made of lossy materials, such as metals.

The variable **lambda** represents the wavelength (in µm) at which the electric field calculation is performed. When computing the excitation electric field, **lambda** must be set to the excitation wavelength; conversely, when calculating the detection PSF, it should correspond to the peak emission wavelength.

To account for the full emission spectrum of a specific fluorophore, the calculations can be repeated for discrete wavelength values across the spectrum. The final result is then obtained as a weighted superposition of these single-wavelength results, using the relative intensities from the emission spectrum as weighting factors.

The variable **focpos** defines the position of the focal plane (in µm) relative to the lowest interface of the bottom multi-layer stack. If **d0** is empty, the focal plane coincides with the bottom interface of the sample layer.

The optional input variable **phirot** allows for the rotation of the input plane wave’s polarization in the BFP. By default, **PlaneWaveExc** calculates the electric fields for an *x*-polarized input plane wave. If **phirot** is specified, the linear excitation polarization is rotated by that angle (**phirot**) with respect to the *x*-axis. As demonstrated in the next chapter, this functionality is sufficient to calculate any arbitrary input polarization structure.

The optional variable **pupil** is the most versatile input, allowing for the definition of arbitrary amplitude and/or phase modulations of the input field in the BFP. If provided, it must be a string describing an executable function of the variables **rad** and **psi**. These variables represent the polar coordinates across the BFP pupil, where **rad** ranges from 0 (at the optical axis) to 1 (at the pupil edge), and **psi** ranges from 0 to 2π. Detailed examples of how to implement the **pupil** variable will be presented in several sections of the following chapter.

Finally, the variable **maxnum** allows for the adjustment of the number of plane wave components used to calculate the electric fields. Internally, the **PlaneWaveExc** function approximates the integral over θ in eq. (4) by summation over discrete values.

By default, the integration region is divided into **maxnum** = 10^3^ equidistant discrete values, for which the integrand is calculated and summed. In certain cases —for instance, when dealing with thick films (several wavelengths in thickness) that exhibit strong forward and backward reflections—this discretization may be too coarse. In such scenarios, **maxnum** can be used to increase the number of discretization points. For all examples discussed in the following chapter, the default value of 10^3^ was sufficient to obtain highly accurate results.

### 2.6 PSF Visualization

The PSF calculation package includes two MATLAB functions designed to visualize computational results. The first function is

**FocusImage3D(xx, yy, zz, field, projectionFac, tsh, clfflag, varargin)** This function renders the PSF as a 3D plot of iso-surfaces at levels of constant intensity. The obligatory input variables are **xx, yy, zz** and **field**. The first three are 3D matrices generated by the **PlaneWaveExc** function if **rhofield** is a structure containing **x** and **y** variables that specify the region of interest. The **field** variable is a 3D array containing the PSF intensity values to be plotted. For an isotropic distribution of emitters, the PSF can be calculated and passed to the function as follows:

**field = abs(exc.fieldx).^2 + abs(exc.fieldy).^2 + abs(exc.fieldz).^2**.

The optional **projectionFac** variable adds maximum intensity projections of the PSF onto the *xy*-, *yz* - and *zx*-planes behind the 3D iso-surface. If it is a single number, then projections are plotted on the surfaces of the current image box enlarged by this number. If **projectionFac = 1**, then projections are plotted directly on the surfaces of the actual image box. If **projectionFac** is a 3D vector, then the image box is enlarged independently in all three dimensions. If **projectionFac** is a 6D vector, then it defines the axis boundaries of the image box on which the projections are plotted. If no projections are desired, **projectionFac** can be omitted or set to an empty matrix [].

The optional **tsh** parameter defines the levels at which iso-surfaces are drawn for the PSF (normalized to its maximum value). Default values are **tsh = 1./exp(1:3)**. This renders three transparent iso-surfaces at 1/*e*, 1/*e*^2^ and 1/*e*^3^ of the PSF maximum value. Generally, **tsh** can be a vector of any length containing values between 0 and 1.

By default, calling **FocusImage3D** invokes the **clf** (clear figure) command. If **clfflag** is provided and not empty, it prevents the function from overriding previous plots. This is particularly useful for overlaying multiple PSFs in a single figure through repeated function calls.

The **varargin** argument allows for passing additional options to modify the plot axes. Internally, these arguments are applied using the command **set(gca, varargin(:))**.

The second visualization function extracts and plots two-dimensional cross sections of the PSF. The calling syntax is:

**[psfcs, x, y] = FocusCrossSection(xx, yy, zz, field, plane, origin, num)**

The input arguments **xx, yy, zz**, and **field** are the same spatial grid and PSF intensity arrays as described for the **FocusImage3D** function. The input variable **plane** is a required 3D vector defined perpendicular to the desired cross-sectional plane (i.e., the plane’s normal vector). If the input variable **origin** is omitted or empty, this plane passes through the coordinate origin (0, 0, 0). If provided as a 3D vector, it defines the specific spatial point through which the cross-section plane passes. An optional two-dimensional vector **num** defines the pixel resolution of the resulting cross-section image. The default resolution is **[200, 200]**.

The function automatically determines the „up” direction for the 2D output: By default, the vertical axis of the cross-section image follows the projection of the global *z*-axis onto the chosen plane. If the cross-section is the *xy*-plane (where the *z*-axis projection would be a null point), the vertical direction defaults to the *y*-axis.

The output variable **psfcs** is a 2D array containing the PSF intensity values for the cross-section, while the **x** and **y** are the coordinate vectors for the horizontal and vertical axes of the 2D image.

The following chapter demonstrates how to apply the **FocusImage3D** and **FocusCrossSection** functions.

## 3. Applications

### 3.1 Focusing through stratified samples: How to deal with refractive index mismatch

As a first and straightforward example, we consider the shape of the intensity distribution when focusing a plane wave through a 1.35 NA oil immersion objective (designed for an immersion medium refractive index of *n*_0_ = 1.51) into a layer of water (*n* = 1.33, thickness *d* = 5 µm) covered by air (*n*_1_ = 1). The excitation wavelength is set to 630 nm. The corresponding MATLAB script for calculating the corresponding excitation PSF images is **PSFModelingFigure3.m**. The computational results are shown in Figure 3, presenting *xz*-cross-sections of the PSF for different focus positions (displacements of the objective’s focal plane measured from the surface of the coverslide).

**Figure 3.**
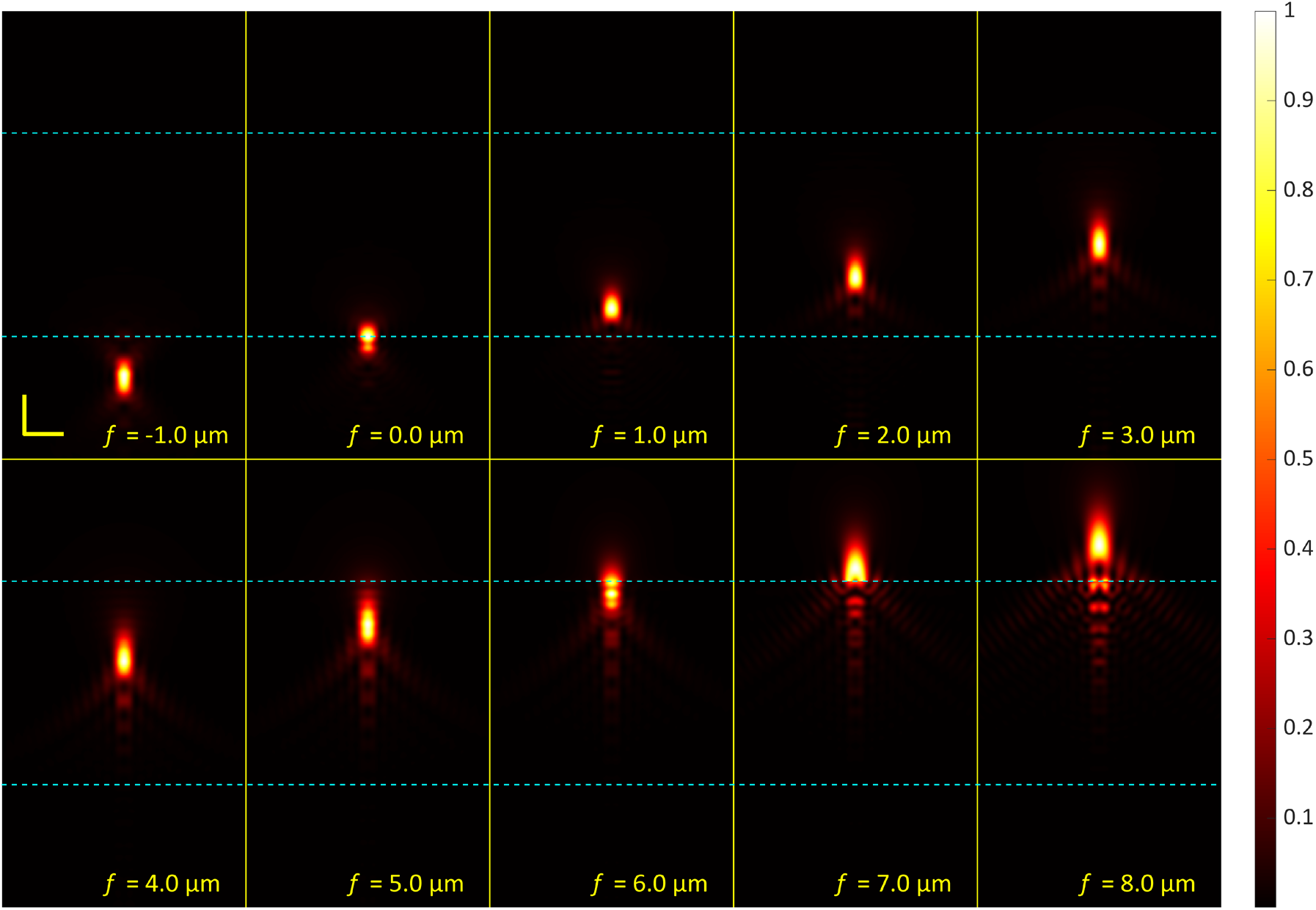
Focusing a plane wave through a 1.35 NA oil-immersion objective into a 5 µm water layer capped by air. The images depict the resulting focal intensity distributions as the focus is translated through the sample, including instances where the nominal focus is located within the glass substrate below or in the air above the water layer. The horizontal lines indicate the positions of the glass/water and water/air interfaces. Note the significant focal distortion and shift caused by the refractive index mismatches between glass (*n* = 1.51), water (*n* = 1.33), and air (*n* = 1.0). The excitation wavelength is 630 nm.

### 3.2 Wide-field microscopy: How to incorporate optical aberrations

Next, we consider the *detection* PSF of a wide-field microscope with aberrations. As was explained in Section 2.4, the detection PSF is nearly identical to the excitation PSF calculated at the emission wavelength, and we keep to this approximation also here. We model the widefield PSF of a microscope equipped with 1.2 NA water immersion objective looking into pure water (*n* = 1.33), assuming an emission wavelength of 670 nm. However, we assume that the imaging optics introduces additional aberrations, which will be described in the conventional way with Zernike polynomials [19].

To incorporate the effect of aberrations, the **pupil** input variable can be utilized. The required phase modulation function is generated by the provided MATLAB routine **Zernike.m**, which encodes the first 36 Zernike polynomials using the Wyant index notation (see https://en.wikipedia.org/wiki/Zernike_polynomials).

To model the aberration with index **j** and having amplitude **amp**, the **pupil** is defined as:

**pupil = [‘exp(2*pi*1i*’ num2str(amp*n/lambda) ‘*Zernike(‘ num2str(j) ‘, rad, psi))’]**

This string encodes the correct pupil function *P* (*u*,ϕ) from eq. (4). Note that within the **PlaneWaveExc** function, the radial variable *u* is denoted as **rad**, and the azimuthal angle ϕ as **psi**.

When modelling detection PSFs, it is important to note that a single call to **PlaneWaveExc** calculates the PSF for only one polarization state (defaulting to *x*-polarized detection). To obtain the full detection profile, the calculation must be repeated by setting the input variable **phirot** to π/2, and the results must then be summed.

Crucially, this rotation also affects the **pupil** function used to describe aberrations. Therefore, for the second call to **PlaneWaveExc**, the **pupil** argument must be modified to:

**pupil = [‘exp(2*pi*1i*’ num2str(amp*n/lambda) ‘*Zernike(‘ num2str(j) ‘, rad, psi+pi/2))’]**

This modification applies a pre-rotation of the pupil function by −π/2 to compensate for the coordinate rotation.

Finally, the default value for the maximum order of the azimuthal expansion, **maxm**, must be increased. Aberrations generate higher-order Fourier modes in the ϕ-expansion of eq. (21), requiring a larger number of **maxm** for convergence. Empirical testing shows that **maxm = 20** is sufficient to yield accurate results for the aberrations shown in Figure 4; further increasing this value does not produce perceptible changes of the output.

**Figure 4.**
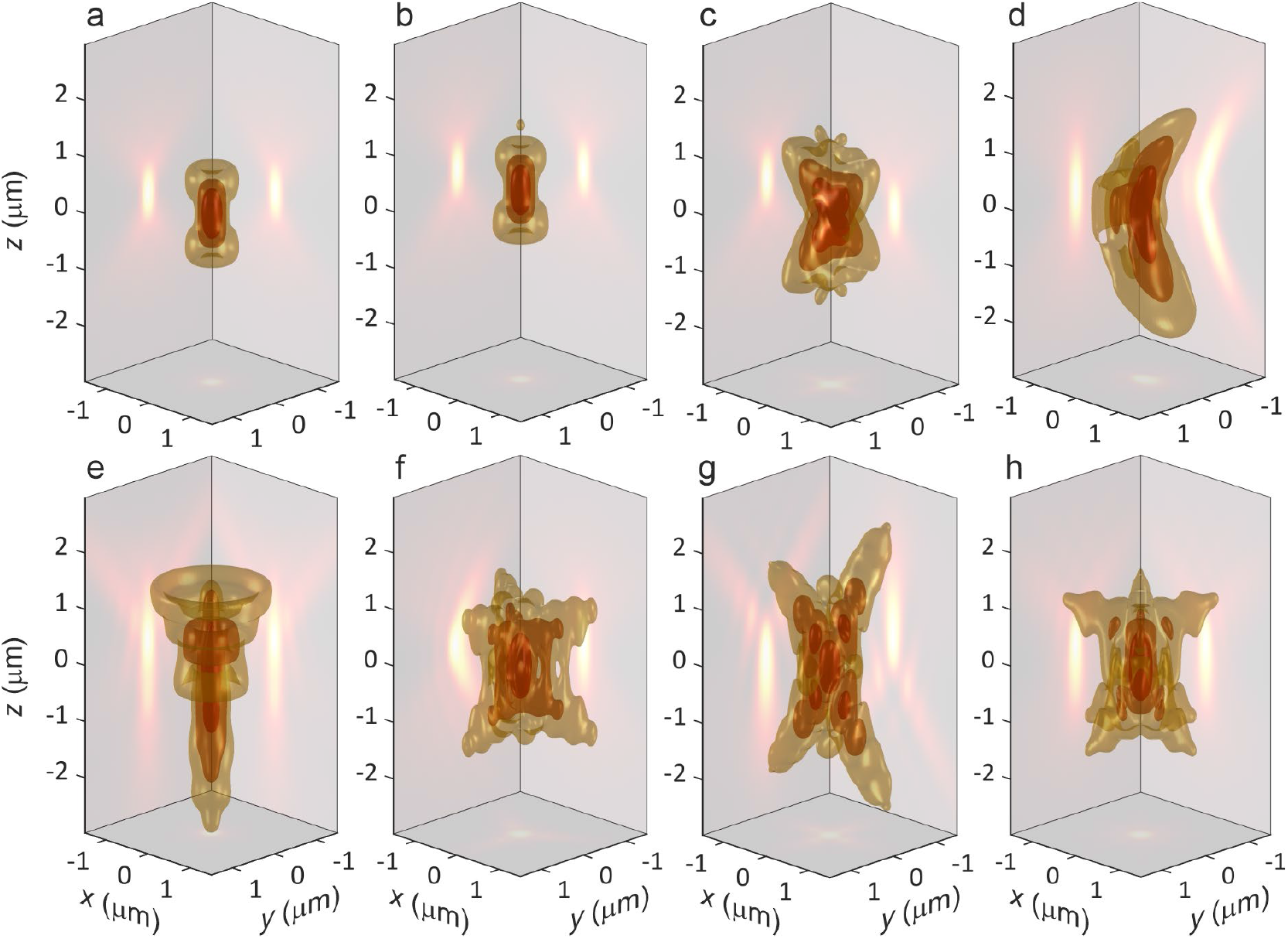
Wide-field PSFs of microscope equipped with a 1.2 NA water immersion objective and for various aberrations. a = no aberrations, b = defocus, c = vertical astigmatism, d = vertical coma, e = primary spherical, f = oblique trefoil, g = vertical secondary astigmatism, h = vertical quadrafoil. The amplitude of the corresponding Zernike polynomials was set to 0.1 of the emission wavelength (i.e. 67 nm) across all simulations. The emission wavelength is 670 nm.

Figure 4 illustrates the computational results for seven prominent aberration types alongside the ideal PSF (panel a). These visualizations were generated using the **FocusImage3D** function, with full implementation details available in the script **PSFModelingFigure4.m**.

### 3.3 Laser Scanning Confocal Microscopy: How to model confocal detection

Modelling the PSF of a Confocal Laser Scanning Microscope (CLSM) [20] requires the combination of two distinct PSFs: the excitation PSF, generated by focusing a plane wave into a diffraction-limited spot, and the confocal detection PSF. The excitation PSF is calculated directly using the **PlaneWaveExc** function. Figure 5a illustrates this result, assuming *x*-polarized excitation light.

**Figure 5.**
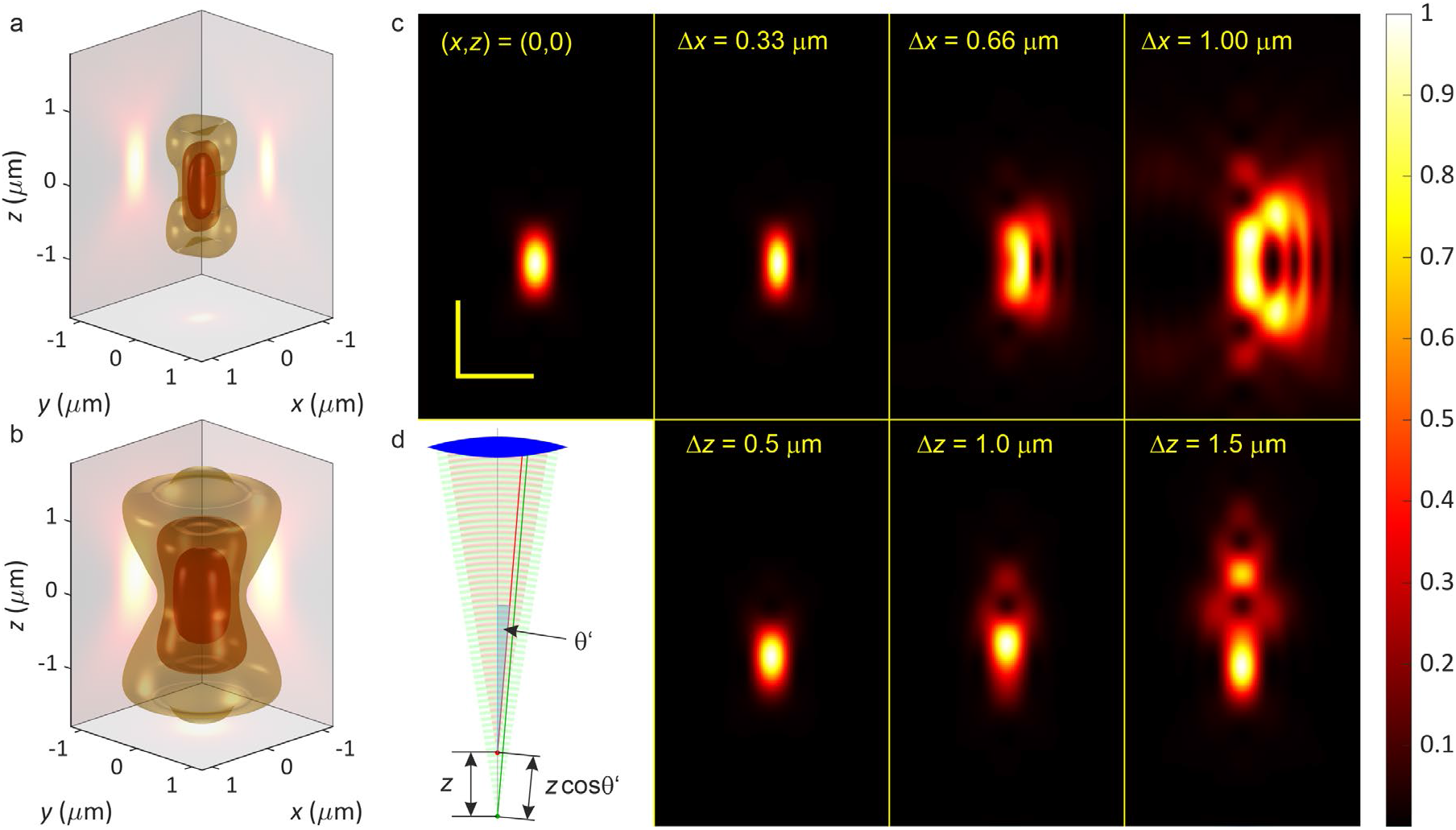
Impact of pinhole misalignment on the PSF of a CLSM. Panel a: Excitation intensity distribution in the focal region for the diffraction-limited focusing of a linearly polarized plane wave. Panel b: The corresponding confocal detection PSF. Panel c: *xz*-cross-sections of the full CLSM PSF. The top row demonstrates the impact of lateral pinhole misalignment (defined by the indicated Δ*x* values), while the bottom row shows the impact of axial misalignment (defined by the indicated Δ*z* values). Computations were done for a 1.2 NA water immersion objective, an excitation wavelength of 630 nm, excitation wavelength of 670 nm, and confocal aperture radius (when back-projected to sample space) of 340 nm. Yellow scale bars represent 1 µm. Panel d: Geometry of phase difference for shifted confocal pinhole plane. When pinhole plane is shifted by Δ*z* towards tube lens, then the optical path of a plane wave traveling along angle θ’ is shorter by Δ*z* cosθ’ as compared to a plane wave coming from the original pinhole plane.

The detection PSF is calculated in two stages: First, the wide-field detection PSF is calculated as described in the previous section (assuming an isotropic emitter dipole orientation). This wide-field result is then convolved with the aperture function, *A*(**ρ**), of a circular pinhole, see eq. (20). The aperture function is defined as:

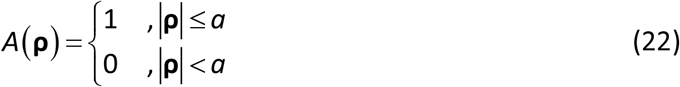

where *a* represents the pinhole radius back-projected into the sample space.

This convolution is performed efficiently using the convolution theorem via Fast Fourier Transforms (FFT). The software package includes a dedicated function, **FourierConvolution.m**, specifically for this purpose. The resulting detection PSF for a perfectly aligned pinhole is displayed in Figure 5b. The total PSF of the CLSM system is determined by the product of the excitation and detection PSFs.

The ideal PSF of the CLSM is displayed in the leftmost panels of Figure 5. These calculations were performed for a 1.2 NA water-immersion objective focusing into water, with an excitation wavelength of 630 nm and an emission wavelength of 670 nm. The pinhole diameter, 2*a*, was set equal to one Airy unit, defined by the position of the first null of the PSF under a scalar approximation (see ref. [21], section III.E). This radius is defined as:

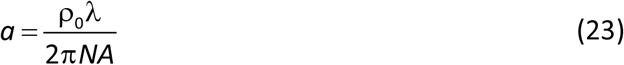

where ρ_0_ is the first zero of the first Bessel function of the first kind, *J*_1_ (ρ), and λ is the emission wavelength. For a wavelength of 670 nm and an NA of 1.2, this back-projected aperture radius in the sample space is 340 nm. Given a system magnification of 60x, this corresponds to a physical pinhole radius of 20.4 µm.

Next, we modelled the impact of pinhole misalignment on the resulting PSF. Modelling lateral displacements is straightforward; the aperture function is modified to *A*(**ρ** − **ρ**_*s*_), where **ρ**_*s*_ represents the lateral shift vector. The resulting PSFs for three different lateral pinhole positions are displayed in the top row of Figure 5c.

To model the effect of axial pinhole displacement, one must consider how the phase relations in the plane wave expansion change when moving the pinhole plane. This is depicted in Figure 5d, which shows that a plane wave traveling at an angle θ’ in image space acquires an extra phase − 2πcosθ′ *z*/λ, where *z* is the shift distance (see also ref. [21], section III.A). Taking into account Abbe’s sine condition, *M* sinθ′ = *n*sinθ (where *M* is the microscopes magnification and *n* the refractive index of the immersion medium), this can be encoded with the **pupil** argument as follows:

**pupil = [‘exp(−1i*’ num2str(2*pi*z/lamem) ‘*cos(asin(‘ num2str(NA/mag) ‘*rad)))’]**

where **mag** is the microscope magnification, and **lamem** is the emission wavelength. Considering that a shift *z* in image space approximately corresponds to a shift of *nz*/*M*^2^ in sample space, we define **z** in the expression above via **z = mag^2/n0*zpin**, where **zpin** is the shift of the confocal aperture back-projected into sample space and **n0** is the refractive index value of water, the objective’s immersion medium, which was equal to the sample’s refractive index value **n**. The corresponding computational results are displayed in the bottom row of Figure 5c.

All computational details for generating the results shown in Figure 5 are contained in the MATLAB script **PSFModellingFigure5.m**.

### 3.4 Defocused imaging of single molecules: How to take into account emission dipole orientation

An important application where the orientation of the fluorescent emitter becomes critical is the defocused wide-field imaging of single molecules with fixed orientation.

Conventionally, this scenario is modelled using a plane wave expansion of the dipole emission field (the Weyl representation) [9,22]. In this framework, each plane wave component is traced through the optical system to the image plane, where all contributions are superimposed to determine the final electromagnetic field distribution. The detected intensity is then calculated as the component of the Poynting vector perpendicular to the detector’s surface, representing the energy flux through that surface.

The principle of reciprocity simplifies this calculation significantly. Instead of tracing emission forward, one calculates the electric field, **E**, generated in the sample space by a single lightemitting pixel on the detector. The detection efficiency for an electric dipole with orientation is then directly proportional to the coupling |**p · E**|^2^.

As a demonstration, we consider the wide-field image of a single molecule in air on a glass coverslide (*n*_*glass*_ = 1.51, *n*_*air*_ = 1.0). Imaging is done with a 1.4 NA oil immersion objective with its immersion medium refractive index perfectly matching that of the glass coverslide. The images are modelled as a function of the molecule’s orientation and the position of the focal plane, using a peak emission wavelength of 670 nm. The computational results are presented in Figure 6, and the implementation details can be found in the accompanying MATLAB script, **PSFModellingFigure6.m**.

**Figure 6.**
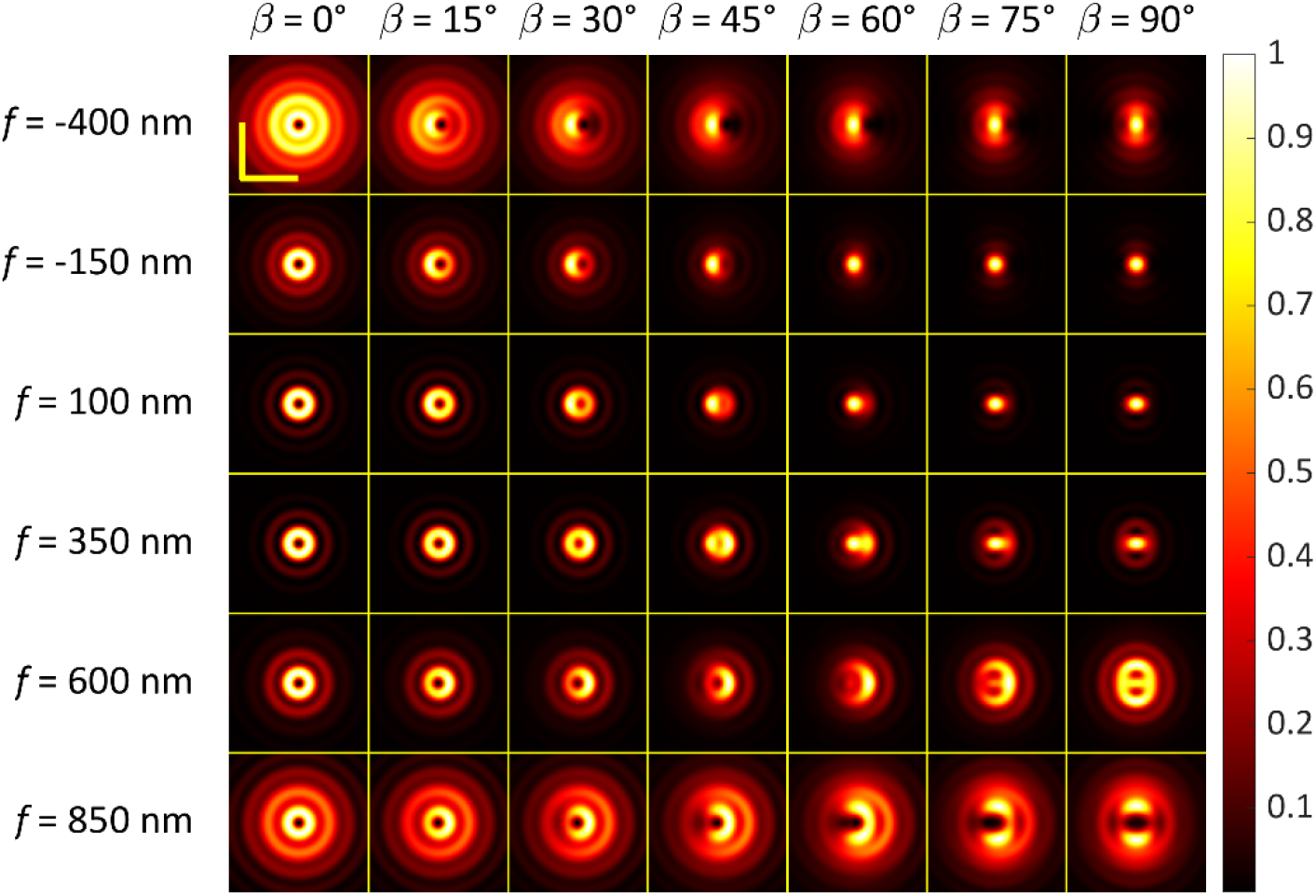
Defocused images of single molecules. Angle β is the inclination between molecule’s emission dipole axis (fixed in the *xz*-plane) and the optical *z*-axis; *f* denotes the defocus value. All panels are normalized to their maximum intensity, with the *x*-axis oriented horizontally. Yellow scale bars: 1 µm. Note that a rotation of the dipole around the optical axis (perpendicular to the image plane) would result in a correspondingly rotated image pattern.

As demonstrated, each molecular orientation corresponds to a unique image pattern, which becomes increasingly distinct as the defocusing distance increases. This relationship allows for the determination of the three-dimensional orientation of single molecules from defocused wide-field images [22–24]—a technique that has found several compelling applications across the fields of chemical physics [25–27] and biology [28].

### 3.5 Scan images of single molecules: How to handle non-trivial excitation polarization

An interesting alternative to wide-field single-molecule imaging is the use of a CLSM [29]. While wide-field imaging primarily probes the orientation of the emission dipole, CLSM instead probes the orientation of the excitation (absorption) dipole. By imaging the same molecule using both techniques, one can infer the relative orientation between the absorption and emission dipoles [30]. Modelling single-molecule images in a CLSM framework is similar to the approach described in the previous section, with the primary difference being that calculations are performed at the excitation wavelength rather than the emission wavelength. For simplicity, we present computational results for a single molecule scanned by a focused excitation beam without a confocal pinhole in the detection pathway. In this configuration, the recorded intensity directly reflects the electric field distribution across the excitation focus as probed by an absorption dipole of a specific orientation. The signal remains proportional to the coupling strength, |**p · E**|^2^, but the structure of the electric field can be significantly more complex than the field distributions used to model the detection PSF in the preceding section.

In this analysis, we consider and compare four distinct focal distributions generated by: (i) a linearly polarized plane wave (the standard configuration in CLSM), (ii) a circularly polarized plane wave, (iii) an azimuthally polarized plane wave—where the electric field vector is oriented tangentially at every point around the optical axis—and (iv) a radially polarized plane wave, where the electric field vector points consistently away from the optical axis. To compute these focal electric field distributions, we utilize the **pupil** function argument to appropriately modulate the phase and vector components of the incident field across the BFP.

Case (i) is straightforward, as the electric field is the direct output of the **PlaneWaveExc** function for an *x*-polarized incident wave. For case (ii), representing circularly polarized excitation, the calculation is repeated with the input parameter **phirot** set to π/2. This second call yields the electric field for a *y*-polarized plane wave; the total complex field is then determined by the superposition of the *x*-polarized result and the *y*-polarized result, the latter multiplied by the imaginary unit *i*.

To generate the focal electric field for azimuthally polarized light, we utilize a two-step superposition. First, **PlaneWaveExc** is called with the pupil function **pupil = ‘-sin(psi)’**. The calculation is then repeated with the same pupil function but with **phirot = pi/2**, which rotates the resulting field by 90° around the optical axis. Summing these two results yields the total focal field for an azimuthally polarized beam. A similar procedure is followed to generate the field for radially polarized light, substituting the pupil function with **pupil = ‘cos(psi)’**.

This method allows for the precise construction of complex vector beams by independently calculating their *x* and *y* contributions. The full implementation of this approach is provided in the accompanying MATLAB script, **PSFModelingFigure7.m**. Finally, single-molecule images are computed by calculating the squared modulus of the scalar product of the molecule’s transition dipole and the electric field distributions.

Figure 7 presents the computational results as a function of the absorption dipole orientation, specifically illustrating the impact of varying inclination angles β relative to the cover glass surface. Consistent with the parameters in the previous section, these simulations were performed for a 1.4 NA oil-immersion objective at an excitation wavelength of 630 nm. The molecule is assumed to be positioned directly on the glass surface, coinciding with the focal plane of the objective.

**Figure 7.**
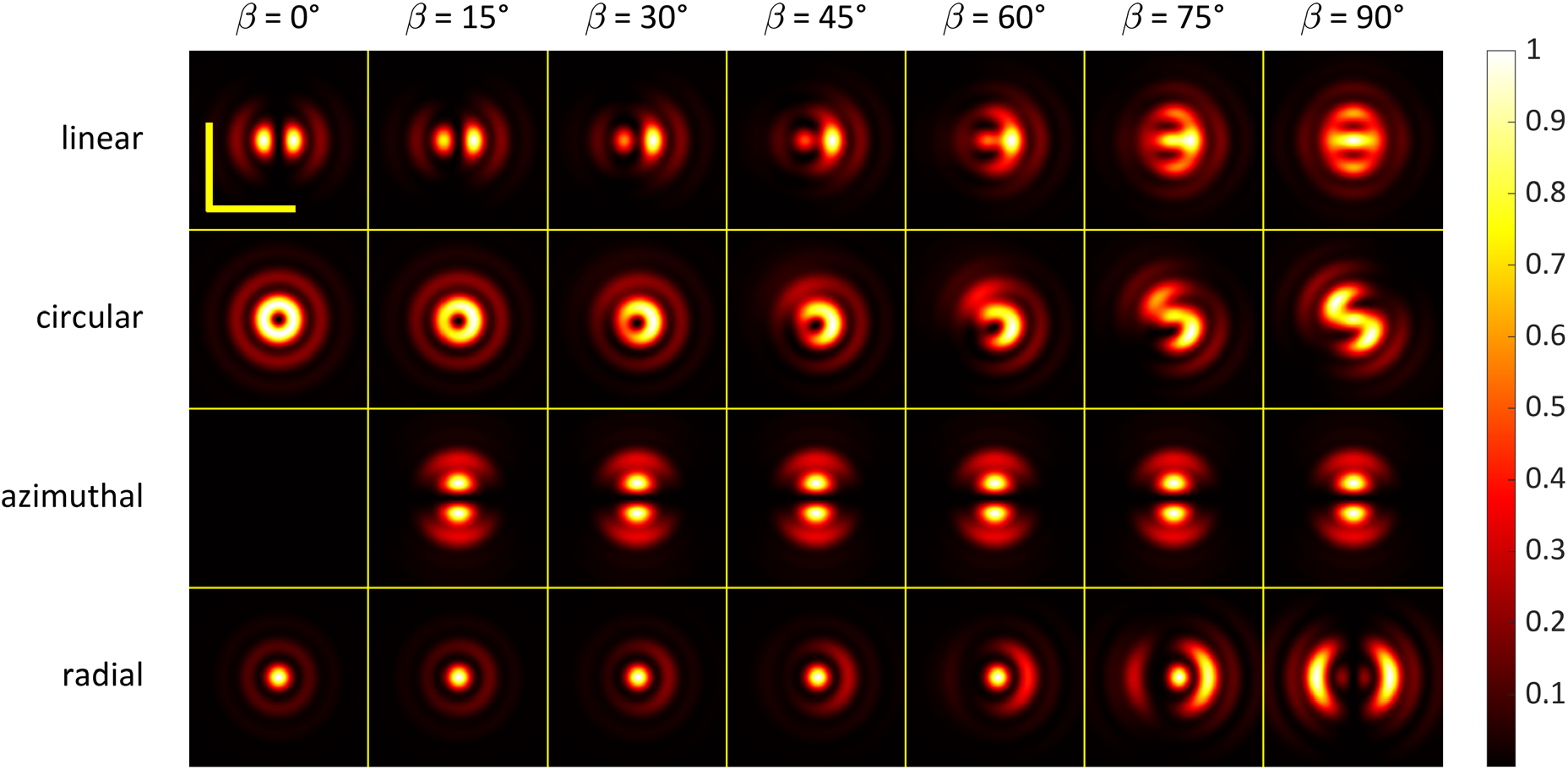
presents scan images of a single molecule for the various excitation focal polarizations indicated to the left of each panel. The polar angle β describes the orientation of the molecule’s absorption dipole relative to the optical axis. An angle β = 0 corresponds to a molecule oriented along the optical axis, while β = 90° represents a molecule lying in the focal plane. The lateral component of the dipole orientation is consistently aligned with the horizontal *x*-axis. All images have been normalized to their respective maximum values. Notably, in the case of azimuthal polarization, no image is generated for β = 0 because this polarization state produces a null electric field along the optical axis. Please note the broken bilateral and mirror symmetry in the images produced with circular excitation polarization. This phenomenon was first reported experimentally in Ref. [31] and is a unique characteristic of focused-beam excitation that could never be observed in conventional wide-field single-molecule imaging. Yellow scale bars are 1 µm.

### 3.6 Stimulated Emission Depletion Microscopy: How to handle complex light fields

Stimulated Emission Depletion (STED) microscopy is a super-resolution technique that by-passes the diffraction limit of light by physically restricting the volume of fluorescence emission [32,33]. It employs two synchronized laser beams: a standard excitation beam to populates the fluorophores’ excited state, and a red-shifted „depletion” beam—often referred to as the STED beam—that forces molecules back to the ground state via stimulated emission before they can spontaneously fluoresce. By shaping the STED beam into a specific geometry with an intensity null (a „hole”) at the centre, only the molecules located at the very heart of the focus are allowed to emit signal, effectively sharpening the PSF beyond the diffraction limit.

Lateral STED is the most common implementation, where a helical phase mask (a vortex ramp from 0 to 2π) is applied to the depletion beam, see Figure 8a. This creates a „doughnut”- shaped focal intensity distribution in the *xy*-plane. Because the intensity of the STED beam increases sharply away from the central null, the lateral periphery of the excitation spot is silenced, resulting in a significantly narrowed lateral PSF.

**Figure 8.**
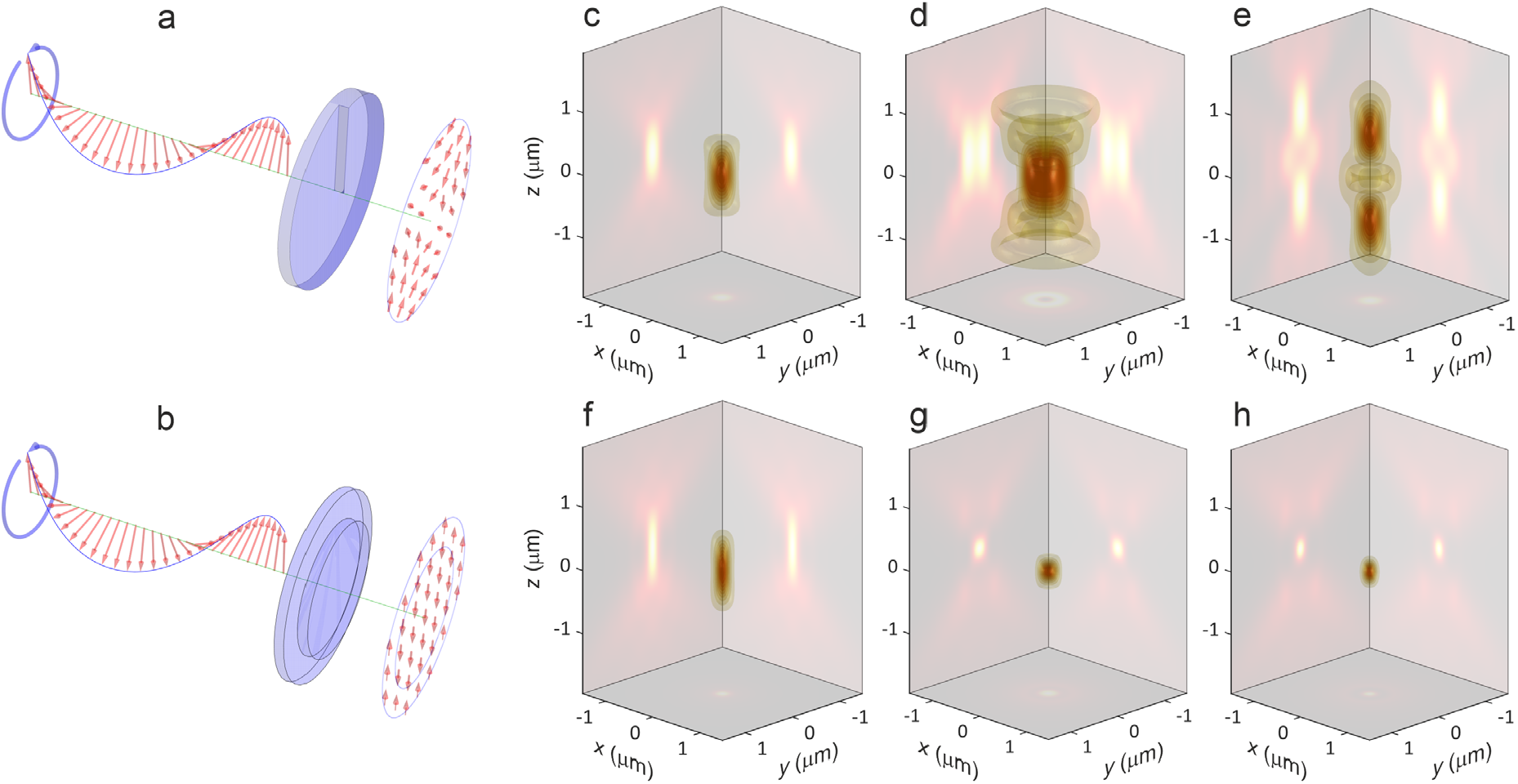
PSF calculations for lateral, axial and isotropic STED. c = fluorescence excitation intensity distribution; d = lateral STED focus generated by helical wave plate shown in panel a, e = axial STED focus generated with the flat-top wave plate shown in panel b; f = lateral STED PSF for a STED intensity of 2 saturation units; g = axial STED PSF for a STED intensity of 10 saturation units; h = STED PSF when using both the lateral and axial STED focus. In the 3D plots, 9 iso-surfaces for evenly spaced values between 0.1 to 0.9 are displayed.

Axial STED improves resolution along the optical axis. To achieve this, a π-step table-top phase mask is applied to the depletion beam, see Figure 8b, creating a „bottle beam” or „double-cap” distribution. This shape places high-intensity depletion zones above and below the focal plane, „sandwiching” the excitation spot and shrinking its axial extent. When lateral and axial depletion patterns are combined, a 3D-STED effect is created.

Modelling the PSF of a STED microscope involves two primary steps: first, calculating the fluorescence excitation focus (which is straightforward) and second, calculating the STED depletion intensity distribution.

Lateral STED is typically generated by focusing an azimuthally polarized beam, a configuration we addressed in the previous section. Axial STED, on the other hand, is created using a flat-top phase plate with a π-phase step. This can be implemented via the pupil function argument **pupil = [‘sign(rad-’ num2str(xi) ‘)’]**, where **xi** (having a value between 0 and 1) defines the radius of the inner phase-shifting disk.

In our modelling of axial STED, as documented in the accompanying script **PSFModelingFigure8.m**, the value of **xi** is determined numerically. First, we compute the central intensity at the focal position for all possible values of **xi** between 0 and 1 and then select the value that results in a central intensity of zero, ensuring a perfect „bottle beam” null. Second, we then use this value in the **pupil** argument for computing the axial STED focus.

Figure 8 presents cross-sections of the fluorescence excitation intensity distribution (panel c), the lateral STED focus (panel d), and the axial STED focus (panel e). The simulations were performed assuming a fluorescence excitation wavelength of 630 nm and a STED wavelength of 700 nm, utilizing a 1.2 NA water-immersion objective focusing into pure water.

To calculate the final STED PSF, we employed the following depletion relation *U*_*STED*_ (**ρ**, *z*) = *U*_*exc*_ (**ρ**, *z*)exp [−ln2 κ*U*_*dep*_ (**ρ**, *z*)], where *U*_*exc*_ is the fluorescence excitation intensity and *U*_*dep*_ is the STED intensity distribution, normalized to its maximum value. The factor κ tunes the relative intensity of the STED beam; a value of κ = 1 signifies that the maximum STED intensity is equal to the saturation intensity for the specific fluorophore used.

Panels f–h present the resulting PSFs for lateral, axial, and combined lateral and axial STED (in all cases, κ was set to 1). These results demonstrate the significant reductions in PSF size along the lateral direction, the axial direction, and both directions simultaneously, highlighting the capability of 3D STED to create a nearly isotropic, sub-diffraction-limited focal volume.

### 3.7 James Webb Space Telescope: How to handle non-trivial amplitude modulations of the pupil functions

As an extreme and highly actual example of amplitude modulation of the pupil function, we consider the image formation in the James Webb Space Telescope (JWST). The JWST is the most powerful space-based observatory ever built, designed to observe the universe in the near- and mid-infrared spectrum [34]. To achieve its unprecedented sensitivity, the telescope utilizes a unique three-mirror anastigmat design that provides a wide field of view while minimizing optical aberrations like coma and astigmatism.

The most striking feature of JWST is its 6.5-meter primary mirror. Because a single mirror of that size would be too large and heavy to fit inside a contemporary rocket fairing, it is constructed from 18 hexagonal segments [34]. These segments are made of gold-plated beryllium, chosen for its extreme stiffness and thermal stability at the cryogenic temperatures (below 50 K) required for infrared astronomy. The hexagonal geometry allows the segments to fold for launch and unfold in space to form a high-fill-factor, nearly circular aperture.

Because the primary mirror is not a continuous circle but a collection of hexagons with gaps and a central obscuration from the secondary mirror, the PSF of the JWST is distinct. The edges of the hexagons and the support struts cause diffraction spikes, resulting in characteristic six-pointed star pattern seen in JWST’s deep-field images. For modelling this system, the „pupil function” must account for this complex hexagonal symmetry and the phase relationships between the 18 individual segments.

We have developed a dedicated function, **jwst_mask_full(rad, psi, a, b)**, which calculates the amplitude modulation of the JWST primary aperture. This function accounts for the hexagonal segmentation of the mirror, the non-reflective gaps of width **a** between segments, and the three support struts of thickness **b**. The function can be used directly as the input for the **pupil** argument. In our simulations, we have set the inter-segment gap **a** to 0.01 and the strut thickness **b** to 0.03, where a value of 1 represents the radius of a disk tightly enveloping the primary mirror.

For the PSF calculations, we have adopted a value of 6.5 m for the primary mirror’s diameter, and a focal length of 131.4 m, which translates into a numerical aperture of NA = 0.0247 for the whole optics. The calculations were done for a wavelength of 1 µm.

Computational results are presented in Figure 9 for the four different amplitude modulation scenarios displayed in the top row. The bottom row shows the resulting PSFs. Due to the low numerical aperture of the optical system, even the PSF for the non-segmented and non-obstructed aperture (Figure 9a, bottom) exhibits a considerable spatial extent and numerous diffraction rings. Our results are in excellent agreement with calculations performed by the NASA JWST team [35]. To accurately calculate these patterns, the **maxm** variable in **PlaneWaveExc** was increased to 200 to account for the presence of high-order angular Fourier modes in the pupil function. The accompanying script, **PSFModeling Figure9.m**, provides the implementation details required to generate the images shown in Figure 9.

**Figure 9.**
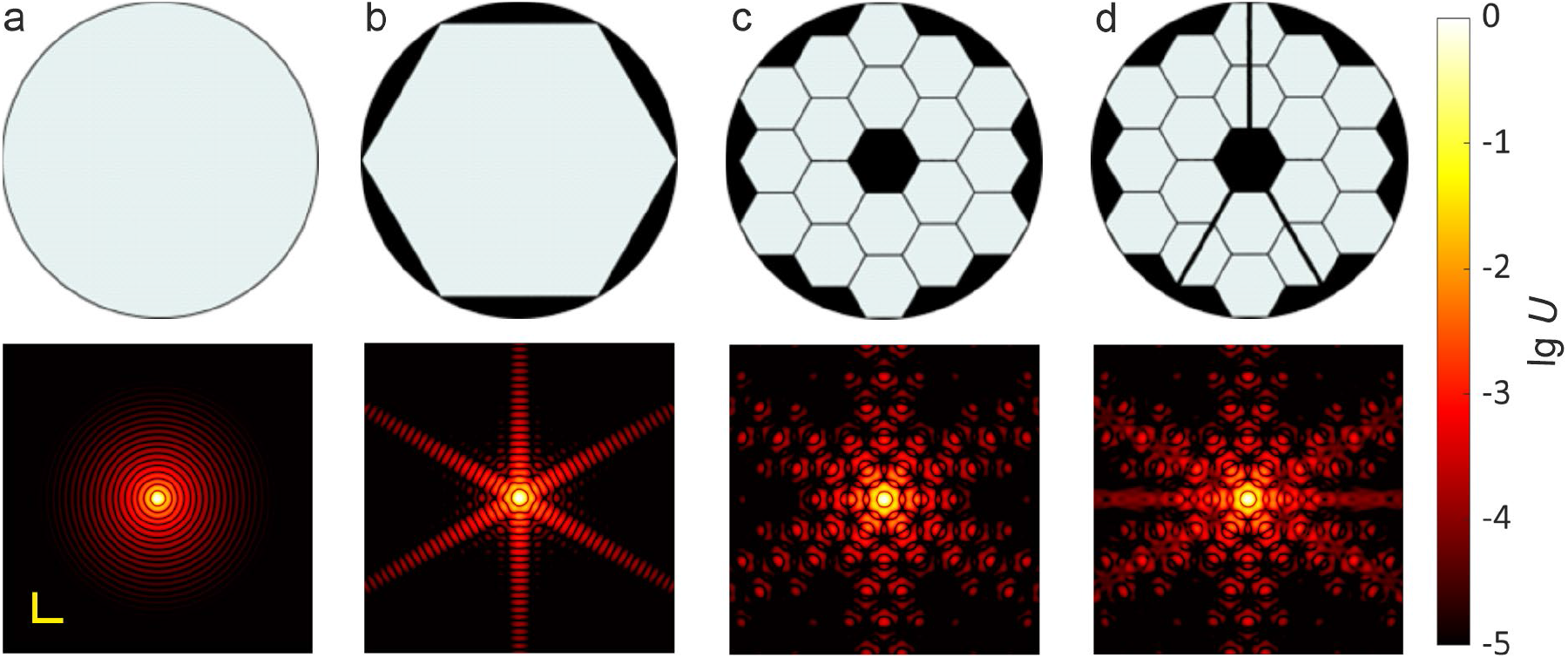
Computational PSF results for various aperture modulations. The top row displays different potential amplitude modulation functions applied over the aperture of the JWST, while the bottom row shows the corresponding PSFs. Panel d provides the closest approximation to the actual geometric configuration of the JWST. Yellow scale bars in the bottom of panel a represent 100 µm (0.1 mm). In contrast to the linear scales used in previous figures, the colour mapping here represents intensity on a logarithmic scale to better visualize the lower-intensity diffraction structures.

### 3.8 Double-helix PSF: How to handle non-trivial phase modulations of the pupil functions

Double-Helix PSF (DH-PSF) microscopy is a powerful wide-field technique designed to extend the axial tracking range and precision of single-molecule localization microscopy. While a standard microscope PSF changes very little when a molecule moves slightly above or below the focal plane—making it difficult to determine the precise *z*-coordinate—the DH-PSF is engineered to rotate its shape based on the emitter’s axial position. By mapping this rotation angle to a specific depth, researchers can reconstruct three-dimensional structures with nanometre precision over a range of several micrometres.

The DH-PSF is created by inserting a specific phase mask into the BFP of the microscope. As a fluorescent molecule moves along the optical axis, the two lobes of the DH-PSF rotate around their common centre. The angular orientation of the line connecting these two lobes serves as a direct proxy for the *z*-position, while the midpoint between the lobes indicates the lateral position.

Initially, Piestun and colleagues introduced the concept [37] using a superposition of Gauss-Laguerre modes [38] for generating the rotating lobes. Subsequently, Prasad proposed an alternative approach using a waveplate composed of Fresnel zones, where successive zones carry spiral phase profiles with increasing topological numbers [36].

In this work, we model the DH detection PSF using the Prasad approach within our pupil function framework. The split Fresnel zone waveplate imprints a pure phase modulation on the wavefront in the BFP, as illustrated in Figure 10a. This action is encoded via the **pupil** argument **pupil = ‘helixPrasad(rad**,**psi)’** where the provided script **helixPrasad.m** calculates the pupil function for the specific phase profile.

**Figure 10.**
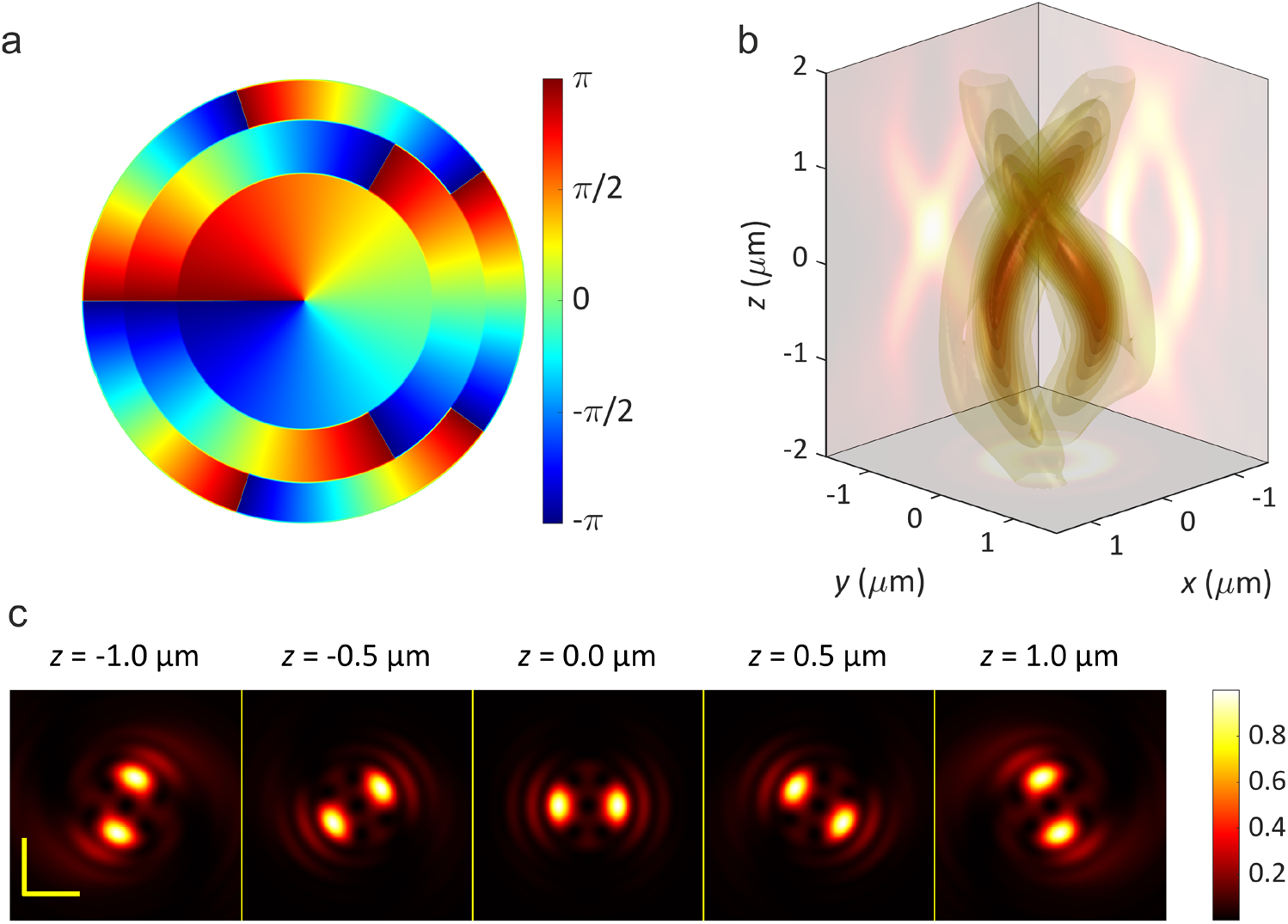
Detection PSF for a Double-Helix (DH) waveplate. Panel a: Displays the imposed phase modulation applied to the incoming wavefront in the BFP, following the design proposed in Ref. [36]. Panel b: Presents the resulting detection PSF as a 3D visualization, where nine distinct intensity isosurfaces are plotted to enhance the visibility of the intricate helical intensity structure. Panel c: Shows five lateral (*xy*) cross-sections of the PSF at the indicated *z*-values, illustrating the rotation of the lobes as a function of axial position.

Consistent with our previous detection PSF calculations, the **PlaneWaveExc** function is called twice—once for *x*-polarized and once for *y*-polarized detection. During the second call, the parameters are set to **phirot = pi/2** and **pupil = ‘helixPrasad(rad, psi+pi/2)’** to account for the coordinate rotation. The resulting detection DH-PSF is presented as a 3D visualization in Figure 10b, with five selected cross-sections along the optical axis displayed in Figure 10c. For these simulations, the emission wavelength was set to 670 nm assuming a 1.2 NA water-immersion objective focusing into water. The generation of these figures is fully documented in the accompanying script, **PSFModelingFigure10.m**.

### 3.9 Super-resolution Optical Fluctuation Imaging: How to deal with non-linearities

An interesting super-resolution microscopy method that only requires a standard wide-field microscope and a fast camera is Super-resolution Optical Fluctuation Imaging or SOFI [39,40]. It is a high-resolution fluorescence microscopy technique that achieves sub-diffraction imaging by analysing the temporal statistics of blinking or fluctuating light emitters. Unlike single-molecule localization techniques that require isolated emitters, SOFI processes a sequence of images where multiple fluorophores may be active simultaneously, provided their brightness fluctuates independently over time. By calculating the *n*^th^ order temporal cumulants of the intensity fluctuations at each pixel, the technique generates a new image where the effective PSF is narrowed by a factor of approximately 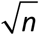, thereby enhancing spatial resolution and contrast while significantly reducing background noise.

The mathematical foundation of SOFI image formation is based on the spatio-temporal distribution of intensity *I* (**ρ**,*t*) on the imaging camera, which can be described as the convolution of the system’s PSF *U*_*det*_ (**ρ** − **ρ**′, *z*′) with a set of *N* point emitters located at positions 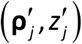. The total intensity at position **ρ** and time *t* is given by

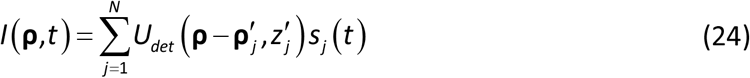

where *s*_*j*_ (*t*) represents the time-dependent brightness of the *j*^th^ emitter. To extract the super-resolution information, SOFI focuses on the intensity fluctuations δ*I* (**ρ**,*t*) = *I* (**ρ**,*t*) − ⟨ *I* (**ρ**,*t*) ⟩ _*t*_. The *n*^th^ order auto-cumulant *C*_*n*_ (**ρ**,τ) of these fluctuations effectively scales the PSF to its *n*^th^ power, 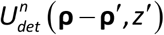. Consequently, the *n*^th^ order SOFI possesses a PSF equivalent to the *n*^th^ power of the original wide-field microscope’s PSF.

This approach is conceptually similar to multi-photon excitation scanning microscopy, with the key difference being that the resulting PSF in multi-photon imaging is a power of the excitation PSF (focal intensity distribution), rather than the detection PSF as in SOFI. Consequently, two-photon excitation scanning microscopy is described by 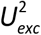, while triple-photon excitation is described by 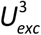. Computational results for SOFI up to the fourth cumulant order are presented in Figure 11 for a wide-field microscope equipped with a 1.2 NA water immersion objective and for an emission wavelength of 670 nm. Full implementation details for the generation of the images in Figure 11 can be found in the accompanying MATLAB script, **PSFModelingFigure11.m**.

**Figure 11.**
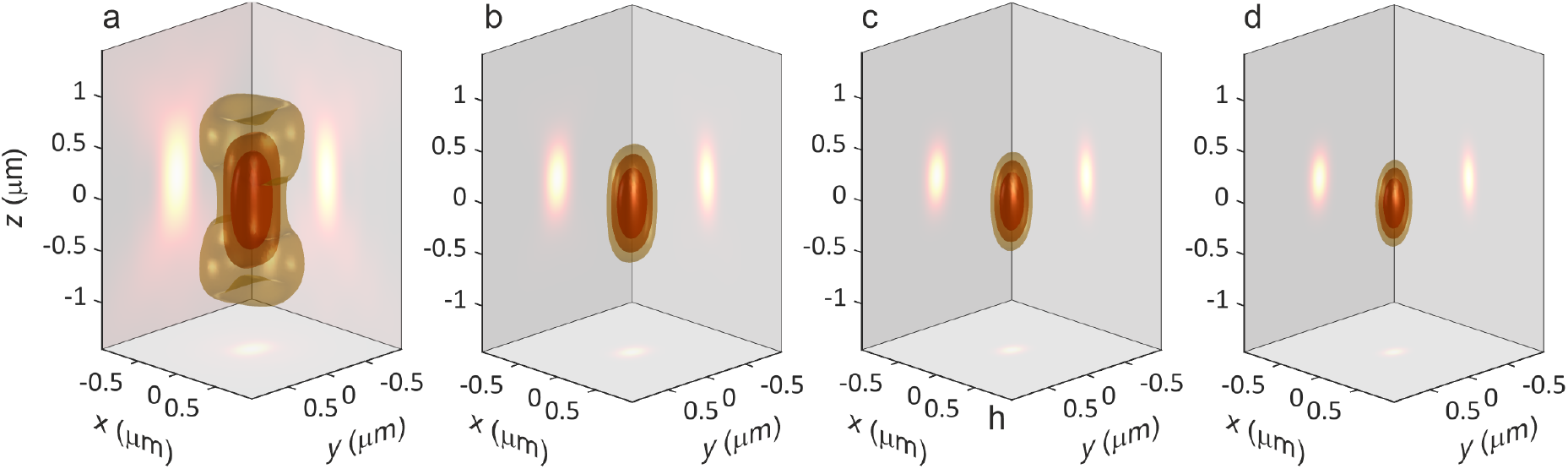
PSF of SOFI up to the fourth order. a = wide-field PSF, b = 2^nd^ order SOFI, c = 3^rd^ order SOFI, d = 4^th^ order SOFI. Please note the reduction of PSF size along all three dimensions. Already 2^nd^ order SOFI provides optical sectioning as in CLSM or two-photon excitation scanning microscopy, in contrast to the wide-field microscope (panel a) used for recording the images from which SOFI is calculated.

### 3.10 iSCAT Microscopy: How to deal with coherent imaging

An important example of coherent, non-fluorescent image formation is interferometric scattering (iSCAT) microscopy [41]. In this modality, the image results from the interference between a reference electromagnetic field (excitation light back-reflected from the glass-water interface) and the electromagnetic field scattered by the sample. In the case of wide-field iSCAT microscopy considered here, the sample is excited with a plane wave traveling along the optical axis, and the back-reflected and scattered fields are imaged onto a camera. In this configuration, the PSF is given by

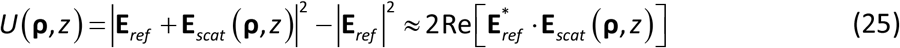

where **E**_*scat*_ (**ρ**, *z*) is the electric field generated at position **ρ** on the camera from a point scatterer at position (0, *z*) in the sample. For the approximation on the r.h.s., we assume that |**E**_*scat*_ (**ρ**, *z*)| ≪ |**E**_*ref*_| at all points.

The incident excitation electric field is modelled as a plane wave traveling along the optical axis, expressed as **E**_*in*_ = **ê**_*x*_ exp [*ik* (*z* − *z*_*f*_)], where *z*_*f*_ is the position of the objectives focal plane and the incident light is assumed to be *x*-polarized. Consequently, the back-reflected reference field on the camera is proportional to **E**_*ref*_ = **ê**_*x*_ · exp [−2*ik* _*f*_], assuming that the glass-water interface is located at *z* = 0.

The scattered field is calculated in two steps: First, the scattering intensity of a point particle is proportional to its electric polarizability, α, multiplied by the incident electric field at the particle’s position. The induced scattering dipole is approximately aligned with the polarization of the incident field (i.e. along the *x*-direction). Second, the electric field generated by this point scatterer on the detector is calculated by using again the principle of reciprocity. Thus, the iSCAT intensity distribution on the detector is

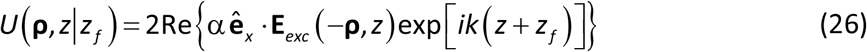

where *z* is the axial position of the scattering particle above the coverslide, and **E**_*exc*_ (**ρ**, *z*) is the electric field distribution generated by an *x*-dipole on the detector at position (**ρ**, *z*) in sample space for a given focal plane position *z*_*f*_. This distribution can be calculated with the **PlaneWaveExc** function.

Figure 12 presents the computational PSFs (PSFs) for wide-field iSCAT microscopy, assuming a point-like gold scatterer. The electric polarizability, α, of such a particle in a medium with refractive index *n* is proportional to the Clausius-Mossotti relation (see Sec. 4.4. in ref. [42])

**Figure 12.**
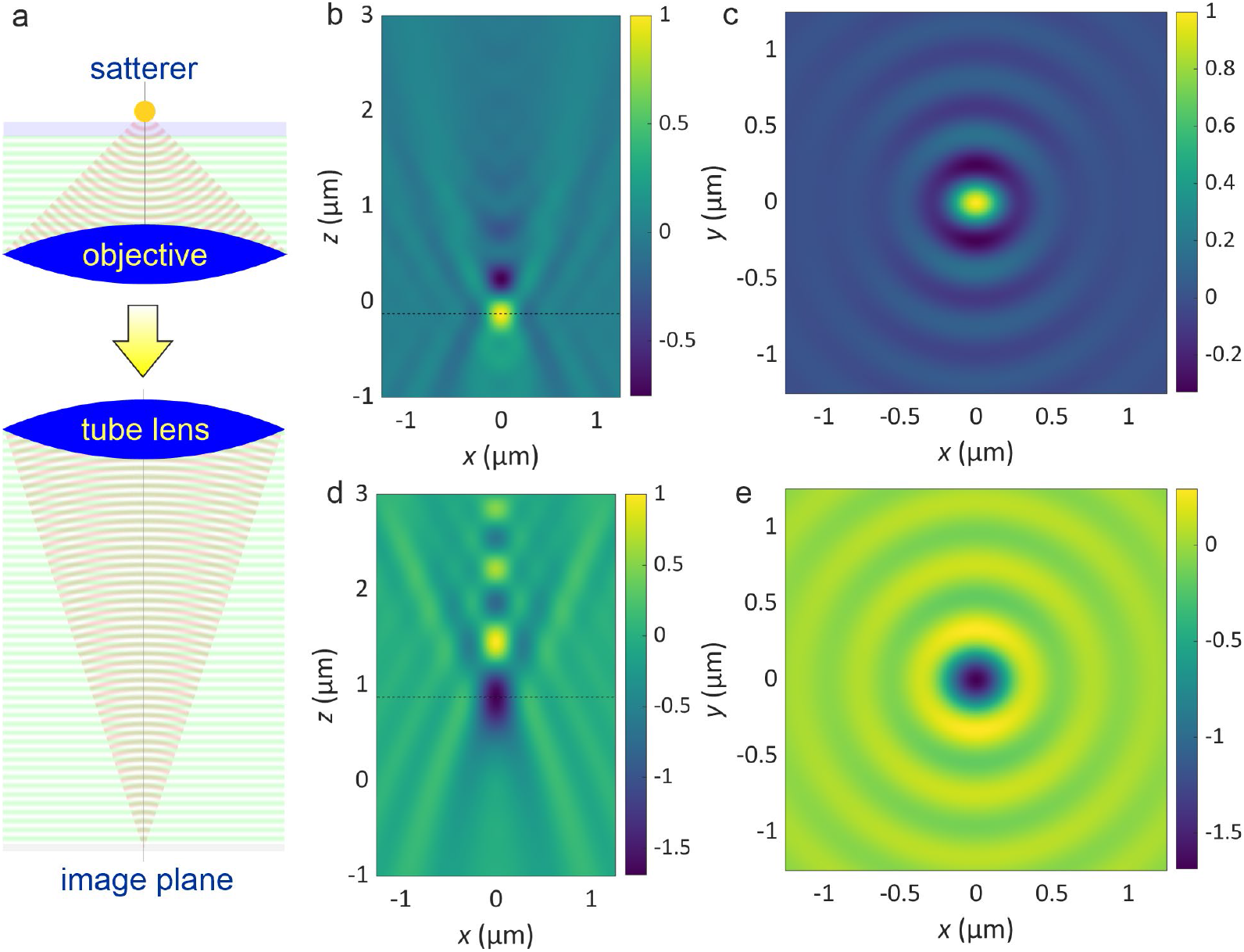
PSF of widefield iSCAT microscopy. Panel a: Schematic of wide-field iSCAT microscopy. Panels b-e: The top row of the figure displays the z-scan PSF of wide-field iSCAT microscopy under optically perfect conditions, where the refractive index of the 170 µm thick coverslip matches the refractive index of the objective’s immersion medium (*n*_*glass*_ = 1.51). In contrast, the bottom row illustrates the resulting *z*-scan PSF when the refractive index of the coverslip deviates by only 0.005 from the ideal value (*n*_*glass*_ = 1.505), a minor mismatch that nonetheless induces perceptible changes in the signal distribution. For both cases, the left panels show the *xz*-cross-sections of the PSF, while the right panels display the *xy*-cross-sections taken at the plane of maximum absolute signal.

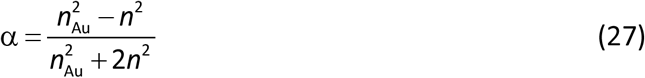

where *n*_Au_ is the complex-valued refractive index of gold. For the selected excitation wavelength of 500 nm, the refractive index of gold was set to *n*_Au_ = 0.8651 + 1.8488 *i*.

The simulation assumes the gold nanoparticle is located directly on the surface of a coverslip (*z* = 0) immersed in pure water (*n*_water_ *=* 1.33). The images depict *z*-scans recorded by sequentially moving the objective’s focal plane along the optical axis; consequently, the images represent (*x, z* _*f*_) cross sections.

The top row of Figure 12 displays cross-sections of the PSF under ideal conditions, where the refractive index of the coverslip perfectly matches that of the objective’s immersion medium. Notably, these ideal conditions failed to reproduce the experimental PSF characteristics reported in Figure 1b and c of reference [43]. A successful reconstruction of the measured PSF was only possible by introducing a slight refractive index mismatch of −0.005 for the 170 µm thick coverslip while keeping the refractive index of the objective’s immersion oil at 1.510, the results of which are shown in the bottom row of Figure 12.

Beyond the axial shift of the focal position (leading to an apparent shift of particle position)— expected due to the differing optical path lengths—the shape of the *z*-scan cross-sections is strikingly different. This divergence stems from the considerable spherical aberration accumulated through the thickness of the coverslip. Our result underscores the extreme sensitivity of interferometric scattering microscopy to even marginal refractive index mismatches, which can profoundly influence final image contrast and symmetry.

Full implementation details for the generation of the images in Figure 12 can be found in the accompanying MATLAB script, **PSFModelingFigure12.m**.

## 4. Conclusion

In this work, we have presented a comprehensive and flexible framework for modelling PSFs across a wide array of microscopy modalities and optical systems. The primary strength of the associated MATLAB package lies in its exceptional „frugality”: the entire suite of applications presented—from multilayered samples to complex phase engineering—relies almost exclusively on one single core function, **PlaneWaveExc**. With a very limited number of input variables, this minimalistic core enables users to handle sophisticated tasks such as incorporating the principle of reciprocity for detection PSF modelling. While this streamlined architecture requires the user to possess a solid understanding of the fundamentals of physical optics to correctly define input parameters—particularly the pupil function for custom modulations—it is precisely this simplicity that makes the package extremely versatile. This approach empowers researchers to model diverse, real-world optical scenarios with high accuracy while maintaining a transparent and adaptable code base.

## 5. Acknowledgments

Ivan Gligonov acknowledges funding by the International Max Planck Research School for Physics of Biological and Complex Systems and by the European Union via the HORIZON–MSCA– 2022–DN „Improving BiomEdical diagnosis through LIGHT-based technologies and machine learning -- BE-LIGHT” (GA number 101119924 -- BE-LIGHT).

Francisco Tenopala-Carmona acknowledges financial support by the Deutsche Forschungsgemeinschaft (DFG, German Research Foundation) through project number 548649522.

Jörg Enderlein acknowledges financial support by the European Research Council (ERC) for financial support via project „smMIET” (GA number 884488) under the European Union’s Horizon 2020 research and innovation program. He also acknowledges financial support by the Deutsche Forschungsgemeinschaft (DFG, German Research Foundation) through project number 565717292, and through Germany’s Excellence Strategy EXC 2067/1-390729940.

Chia-Lung Hsieh and Jörg Enderlein thank Mr. Yan-Hsien Chen for fruitful discussions of iSCAT microscopy.

## References

[1] D. A. Agard and J. W. Sedat, Three-dimensional architecture of a polytene nucleus, Nature 302, 5910 (1983).

[2] D. S. C. Biggs, 3D Deconvolution Microscopy, Curr. Protoc. Cytom. 52, 12.19.1 (2010).

[3] M. Leutenegger, R. Rao, R. A. Leitgeb, and T. Lasser, Fast focus field calculations, Opt. Express 14, 11277 (2006).

[4] J. Lin, O. G. Rodríguez-Herrera, F. Kenny, D. Lara, and J. C. Dainty, Fast vectorial calculation of the volumetric focused field distribution by using a three-dimensional Fourier transform, Opt. Express 20, 1060 (2012).

[5] M. Hillenbrand, A. Hoffmann, D. P. Kelly, and S. Sinzinger, Fast nonparaxial scalar focal field calculations, JOSA A 31, 1206 (2014).

[6] J. Kim, Y. Wang, and X. Zhang, Calculation of vectorial diffraction in optical systems, JOSA A 35, 526 (2018).

[7] R. Tong, Z. Dong, Y. Chen, F. Wang, Y. Cai, and T. Setala, Fast calculation of tightly focused random electromagnetic beams: controlling the focal field by spatial coherence, Opt. Express 28, 9713 (2020).

[8] L. Mandel and E. Wolf, Optical Coherence and Quantum Optics (Cambridge University Press, Cambridge, 1995).

[9] L. H. Novotny Bert, Principles of Nano-Optics: Resolution and Locali ation, in Principles of Nano-Optics, Vol. NA (2012), pp. 86–130.

[10] G. Kirchhoff, I. On the relation between the radiating and absorbing powers of different bodies for light and heat, Lond. Edinb. Dublin Philos. Mag. J. Sci. 20, 1 (1860).

[11] R. Carminati, M. Nieto-Vesperinas, and J.-J. Greffet, Reciprocity of evanescent electro-magnetic waves, J. Opt. Soc. Am. A 15, 706 (1998).

[12] R. Carminati, J. J. Sáenz, J.-J. Greffet, and M. Nieto-Vesperinas, Reciprocity, unitarity, and time-reversal symmetry of the S matrix of fields containing evanescent components, Phys. Rev. A 62, 012712 (2000).

[13] R. J. Potton, Reciprocity in optics, Rep. Prog. Phys. 67, 717 (2004).

[14] H. Benisty, J.-J. Greffet, and P. Lalanne, Introduction to Nanophotonics (Oxford university press, 2022).

[15] E. Wolf, Electromagnetic diffraction in optical systems-I. An integral representation of the image field, Proc. R. Soc. Lond. Ser. Math. Phys. Sci. 253, 349 (1959).

[16] Richards B. and Wolf E., Electromagnetic diffraction in optical systems, II. Structure of the image field in an aplanatic system, Proc. R. Soc. Lond. Ser. Math. Phys. Sci. 253, 358 (1959).

[17] M. Born and E. Wolf, Principles of Optics: Electromagnetic Theory of Propagation, lnterference and Diffraction of Light, 7th ed. (Cambridge University Press, Cambridge, 1999).

[18] I. Gregor, D. Patra, and J. Enderlein, Optical Saturation in Fluorescence Correlation Spectroscopy under Continuous-Wave and Pulsed Excitation, ChemPhysChem 6, 164 (2005).

[19] I. Gligonov and J. Enderlein, Variational calculus approach to Zernike polynomials with application to FCS, Biophys. J. 124, 3366 (2025).

[20] M. Minsky, Memoir on inventing the confocal scanning microscope, Scanning 10, 128 (1988).

[21] M. Fazel, K. S. Grussmayer, B. Ferdman, A. Radenovic, Y. Shechtman, J. Enderlein, and S. Pressé, Fluorescence microscopy: A statistics-optics perspective, Rev. Mod. Phys. 96, 025003 (2024).

[22] M. Btihmer and J. Enderlein, Orientation imaging of single molecules by wide-field epifluorescence microscopy, J. Opt. Soc. Am. B 20, 554 (2003).

[23] D. Patra, I. Gregor, and J. Enderlein, Image Analysis of Defocused Single-Molecule Images for Three-Dimensional Molecule Orientation Studies, J. Phys. Chem. A 108, 6836 (2004).

[24] D. Patra, I. Gregor, J. Enderlein, and M. Sauer, Defocused imaging of quantum-dot angular distribution of radiation, Appl. Phys. Lett. 87, 101103 (2005).

[25] H. Uji-i, S. M. Melnikov, A. Deres, G. Bergamini, F. De Schryver, A. Herrmann, K. Müllen, J. Enderlein, and J. Hofkens, Visualizing spatial and temporal heterogeneity of single molecule rotational diffusion in a glassy polymer by defocused wide-field imaging, Polymer 47, 2511 (2006).

[26] P. Dedecker, B. Muls, A. Deres, H. Uji-i, J. Hotta, M. Sliwa, J.-P. Soumillion, K. Müllen, J. Enderlein, and J. Hofkens, Defocused Wide-field Imaging Unravels Structural and Temporal Heterogeneity in Complex Systems, Adv. Mater. 21, 1079 (2009).

[27] A. Deres, G. A. Floudas, K. Müllen, M. Van der Auweraer, F. De Schryver, J. Enderlein, H. Uji-i, and J. Hofkens, The Origin of Heterogeneity of Polymer Dynamics near the Glass Temperature As Probed by Defocused Imaging, Macromolecules 44, 9703 (2011).

[28] E. Toprak, J. Enderlein, S. Syed, S. A. McKinney, R. G. Petschek, T. Ha, Y. E. Goldman, and P. R. Selvin, Defocused orientation and position imaging (DOPI) of myosin V, Proc. Natl. Acad. Sci. 103, 6495 (2006).

[29] J. Dapprich, O. Mets, W. Simm, M. Eigen, and R. Rigler, Confocal scanning of single molecules, Exp Tech Phys 41, 259 (1995).

[30] N. Karedla, S. C. Stein, D. Hähnel, I. Gregor, A. Chizhik, and J. Enderlein, Simultaneous Measurement of the Three-Dimensional Orientation of Excitation and Emission Dipoles, Phys. Rev. Lett. 115, 173002 (2015).

[31] D. Marx, I. Gligonov, D. Malsbenden, D. Wtill, O. Nevskyi, and J. Enderlein, Mapping Optical Chirality with Single Fluorescent Molecules, Nano Lett. (2026).

[32] S. W. Hell and J. Wichmann, Breaking the diffraction resolution limit by stimulated emission: stimulated-emission-depletion fluorescence microscopy, Opt. Lett. 19, 780 (1994).

[33] T. A. Klar, S. Jakobs, M. Dyba, A. Egner, and S. W. Hell, Fluorescence microscopy with diffraction resolution barrier broken by stimulated emission, Proc. Natl. Acad. Sci. 97, 8206 (2000).

[34] J. P. Gardner et al., The James Webb Space Telescope, Space Sci. Rev. 123, 485 (2006).

[35] R. B. Makidon, S. Casertano, C. Cox, and R. van der Marel, The JWST point spread function: Calculation methods and expected properties, Tech. Rep. JWST-STScI-001157 1157 (2007).

[36] S. Prasad, Rotating point spread function via pupil-phase engineering, Opt. Lett. 38, 585 (2013).

[37] G. Grover, K. DeLuca, S. Quirin, J. DeLuca, and R. Piestun, Super-resolution photon-efficient imaging by nanometric double-helix point spread function localization of emitters (SPINDLE), Opt. Express 20, 26681 (2012).

[38] J. Enderlein and F. Pampaloni, Unified operator approach for deriving Hermite–Gaussian and Laguerre–Gaussian laser modes, J. Opt. Soc. Am. A 21, 1553 (2004).

[39] T. Dertinger, R. Colyer, G. Iyer, S. Weiss, and J. Enderlein, Fast, background-free, 3D super-resolution optical fluctuation imaging (SOFI), Proc. Natl. Acad. Sci. 106, 22287 (2009).

[40] T. Dertinger, J. Xu, O. Naini, R. Vogel, and S. Weiss, SOFI-based 3D superresolution sectioning with a widefield microscope, Opt. Nanoscopy 1, 2 (2012).

[41] R. G. Mahmoodabadi, R. W. Taylor, M. Kaller, S. Spindler, M. Mazaheri, K. Kasaian, and V. Sandoghdar, Point spread function in interferometric scattering microscopy (iSCAT). Part I: aberrations in defocusing and axial localization, Opt. Express 28, 25969 (2020).

[42] J. D. Jackson, Classical Electrodynamics, 3rd ed. (Wiley-VCH, 2021).

[43] M. Küppers, D. Albrecht, A. D. Kashkanova, J. Lühr, and V. Sandoghdar, Confocal interferometric scattering microscopy reveals 3D nanoscopic structure and dynamics in live cells, Nat. Commun. 14, 1962 (2023).

